# Nanobody Mediated Macromolecular Crowding Induces Membrane Fission and Remodeling in the African Trypanosome

**DOI:** 10.1101/2021.01.13.426364

**Authors:** Alexander Hempelmann, Laura Hartleb, Monique van Straaten, Hamidreza Hashemi, Johan P. Zeelen, F. Nina Papavasiliou, Markus Engstler, C. Erec Stebbins, Nicola G. Jones

**Affiliations:** Division of Structural Biology of Infection and Immunity, German Cancer Research Center, Heidelberg, Germany; Department of Cell and Developmental Biology, Theodor-Boveri-Institute, Biocenter, University of Würzburg, Würzburg, Germany; Division of Immune Diversity, German Cancer Research Center, Heidelberg, Germany

**Keywords:** African Trypanosome, Host-Pathogen Interaction, Variant Surface Glycoproteins, Immune Epitope Mapping, Structural Biology, Nanovesicle Formation, Nanotube Formation, Protein Crowding, Membrane Fission

## Abstract

The dense Variant Surface Glycoprotein (VSG) coat of African trypanosomes represents the primary host-pathogen interface. Antigenic variation prevents clearing of the pathogen by employing a large repertoire of antigenically distinct VSG genes, thus neutralizing the host’s antibody response. To explore the epitope space of VSGs, we generated anti-VSG nanobodies and combined high-resolution structural analysis of VSG-nanobody complexes with binding assays on living cells, revealing that these camelid antibodies bind deeply inside the coat. One nanobody caused rapid loss of cellular motility, possibly due to blockage of VSG mobility on the coat, whose rapid endo-and exocytosis is mechanistically linked to *T. brucei* propulsion and whose density is required for survival. Electron microscopy studies demonstrated that this loss of motility was accompanied by rapid formation and shedding of nanovesicles and nanotubes, suggesting that increased protein crowding on the dense membrane can be a driving force for membrane fission in living cells.

## Introduction

*Trypanosoma brucei* and related parasites cause African sleeping sickness in humans and Nagana in cattle. A distinguishing feature of these pathogens is a densely packed surface coat that is populated almost entirely by the **V**ariant **S**urface **G**lycoprotein (**VSG**) in the mammalian-infectious forms of the parasite. As *T. brucei* is an extracellular pathogen, the VSG coat is extensively exposed to the immune system and is the primary surface of interaction within the host. It is also highly antigenic, leading quickly to robust antibody mediated responses that are neutralizing (Borst, 2002; Overath et al., 1994). However, parasites possess an extensive genetic repertoire of antigenically distinct VSGs (2000+ genes and pseudogenes), allowing individual parasites to switch the coat to another variant, thus creating a novel surface array to which the immune system is naïve and must readapt. This process of antigenic variation repeats with recurring cycles of parasite clearance, coat switching, and pathogen regrowth (Bangs, 2018).

In addition to antigenic variation, trypanosomes employ another immune evasion mechanism, antibody clearance and subsequent recycling of VSG molecules. This process helps evasion of complement-mediated immune destruction but is only effective at low antibody titer (Engstler et al., 2007; Horn, 2014; Mugnier et al., 2016; Pal et al., 2003). Facilitated by the remarkably high endocytosis rate at the flagellar pocket, anti-VSG antibodies are selectively removed from VSG molecules within minutes (Engstler et al., 2004; Webster et al., 1990). This requires high mobility of the VSG molecules and free lateral diffusion (Bülow et al., 1988; Hartel et al., 2016, 2015).

The VSGs are glycosylated, homodimeric proteins of roughly 60 kDa comprised of two subdomains: a long, rod-like N-terminal domain (NTD) spanning approximately 350 amino acids followed by a C-terminal domain (CTD) of roughly 100 amino acids that attaches to the glycophosphatidylinositol (GPI) anchor which links the protein to the membrane. The antigenic region of the protein is considered to be the NTD and, in particular, the upper regions of the NTD that would be exposed to the immune system in the dense packing of tens of millions of VSG proteins on the surface (Schwede et al., 2015). The NTDs are composed of a tightly packed three-helix-bundle decorated with extended insertions that produce top and bottom “lobes” at polar ends of the rod-like, helical scaffolding.

Despite decades of research, very little is known about the interactions of immune proteins with the VSG coat. Few studies have mapped any defined natural epitopes and at present no antibody-VSG co-crystal structures exist. Reviewing multiple indirect studies (involving monoclonal antibodies, nanobodies, polyclonal antisera, lectins, and proteases), Schwede et al. propose that contrary to many discussions in published literature, a substantial portion of the N-terminal domain (ending at the bottom lobe) of the VSGs could be accessible to the immune system (Schwede et al., 2015).

In order to address this topic, we pursued structural studies of several camelid nanobodies raised to the VSG coat. These provide both a biologically relevant context (as *T. brucei* related species cause surra in camelids – (Radwanska et al., 2018)), as well as a system generally amenable to crystallographic analysis (de Marco, 2011). While camels and llamas have a very similar immune system to humans and rodents, there are variations. Nanobodies (Nbs) are single chain, minimal variable region-containing antibody fragments consisting of one protein domain. They tend to be highly stable and to crystallize very well and thus are useful alternatives to studying pathogen-immune interactions if the larger mouse and human antibodies are challenging to crystallize.

Previous studies have shown that high-affinity Nbs can be effectively raised to VSGs (Arbabi-Ghahroudi, 2017). Addition of such Nbs to AnTat1.1 expressing *T. brucei* was shown to induce trypanosome lysis even in the absence of the effector Fc domain (Stijlemans et al., 2017). This effect was described to initiate quick immobilization of trypanosomes, enlargement of the flagellar pocket and a major blockage of endocytosis. However, the mechanism mediating these phenomena have remained obscure.

In this study, we used the camelid immune system (Figure S1) to generate a series of nanobodies to *T. brucei* VSG2 (also known as VSG221 or VSG MITat1.2). X-ray crystallographic studies reveal that these nanobodies bind not to the upper, exposed surface of the elongated VSG molecules, but instead as much as 100 Å from the top. Indeed, we find that these nanobodies and their corresponding camelid antibodies are also able to recognize the intact trypanosome coat on living cells. These results demonstrate that VSG epitopes buried deeply in the dense surface coat are in fact accessible to B-cells and secreted antibodies.

Similar to prior reports, this binding can be highly toxic to the parasite (Stijlemans et al., 2011, 2017) and we observed paralysis of the normally highly mobile trypanosomes within minutes of nanobody addition. This paralysis was linked to a nanobody induced block on VSG diffusion in the coat. Parasite movement and endo-and exocytosis of the coat are mechanistically linked and essential for survival in the mammalian host. At the same time, VSG density on the coat is highly regulated, and significant raising of the density adversely impacts VSG movement in the membrane (Hartel et al., 2016), making the African trypanosome a unique model system for studying the effects of protein crowding in living cells. Examination of the parasite by electron microscopy after Nb binding demonstrated that loss of VSG mobility (and subsequent loss of cellular motility) was accompanied by rapid formation and shedding of nanovesicles and nanotubes, suggesting that increased protein crowding on this already dense membrane physically drives membrane fission – the first example of this phenomenon in living cells.

## Results

### Production and Crystal Structures of VSG2-Nanobody Complexes

Nanobodies were generated from llamas immunized with the N-terminal domain of VSG2 (Methods). To ensure that the cloned genes were able to bind within surface coats and not only the soluble form of the VSG, a filtering step of panning the library of B-cell clones against intact UV-inactivated trypanosomes was added and all clones were scored by flow cytometry analysis as binding to intact, live trypanosomes. From this we obtained a library of 69 distinct nanobodies (belonging to 30 different CDR3 groups or B-cell lineages). We initially crystallized three different clones complexed to VSG2 (NB9_VSG2_, NB11_VSG2_, and NB14_VSG2_, Figure 1, Methods, Figure S2, Table S1 and S2). Those structures revealed that all the nanobodies bound similarly to the same epitope region and possessed a highly conserved consensus set of amino acids in the CDR1 and CDR3 loops that characterized the majority of clones in the library, despite being derived from independent B-cell lineages (discussed below). To identify other binding modalities, we searched for clones in our library with the most divergent CDR regions. A few such were identified, and we crystallized one of these (NB19_VSG2_) which indeed bound in a distinct location on VSG2. However, unexpectedly, this was not on the top, putative antigenic surface of the molecule, but nearly 100 Å down from the upper surface at the bottom of the N-terminal domain and adjacent to the postulated location of the CTD (Bartossek et al., 2017). The overall structure of VSG2 is not significantly changed from that previously reported (Freymann et al., 1990), aligning with a root-mean-square-deviation in 1440 atomic positions of 0.5 Å (single monomer).

**Figure 1.**
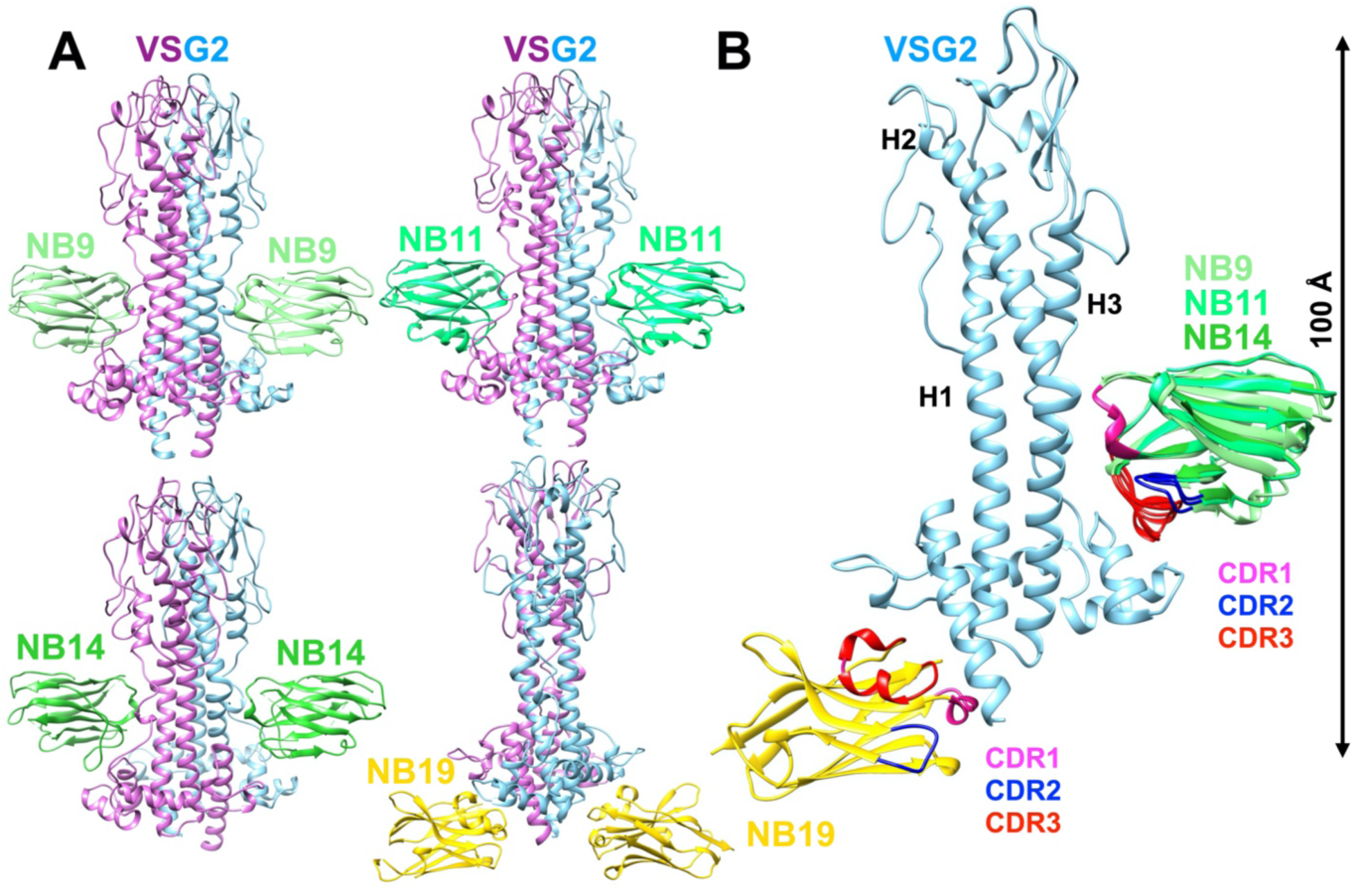
Overall Structures of VSG2-NB Complexes. **(A)** The four nanobody-VSG2 structures with NB9 (NB9 in light green, top left), NB11 (NB11 in lime green, top right), NB14 (NB14 in green, bottom left) and NB19 (NB19 in gold, bottom right), and VSG2 (dimer in cyan/purple) are depicted as ribbon diagrams. Superposition of VSG2-NB complexes. The Complementarity-Determining Regions (CDR) are shown in magenta (CDR1), blue (CDR2) and red (CDR3). Structures illustrated with Chimera (Pettersen et al., 2004).

### Nanobodies Bind Deep into the VSG Coat

Three of the nanobodies (NB9_VSG2_, NB11_VSG2_, and NB14_VSG2_) bind in an identical position. The binding site is approximately 60 Å down from the upper lobe of VSG2 and formed by a concave surface on the VSG 3-helix bundle about midway down the length of the rod-like NTD (Figure 1A and B). This is consistent with a hypothesis that nanobodies are more likely to bind concave or convex surfaces than flat ones (Stijlemans et al., 2004). Numerous side chains from the second two helices of the helical bundle make contacts to the nanobodies (residues spanning 100-115 and 280-302 of VSG2), with the primary contacts solvent-facing, predominantly hydrophilic residues (e.g. Q111, T112, A285, E286, T301, and D313) as well as several minor side chain contacts and backbone contacts (Figure 2A). These nanobodies interact with VSG2 through their variable CDR1 and CDR3 loops (without any significant contacts from the CDR2 loop) with a consensus set of side chains (CDR1 residues T28, S30, N31 and CDR3 residues L100, L101, S103, Figure 2, A and C). In contrast, NB19_VSG2_ binds underneath the bottom lobe of VSG2 purely through CDR3 (Figure 2B). The contacts are a mixture of hydrophobic and polar interactions, but with a preponderance of aromatic side chain stacking (π-stacking) interactions involving residues W102, W112, and Y115 of NB19_VSG2_ with the side chains Y375 and Y376 of VSG2.

**Figure 2.**
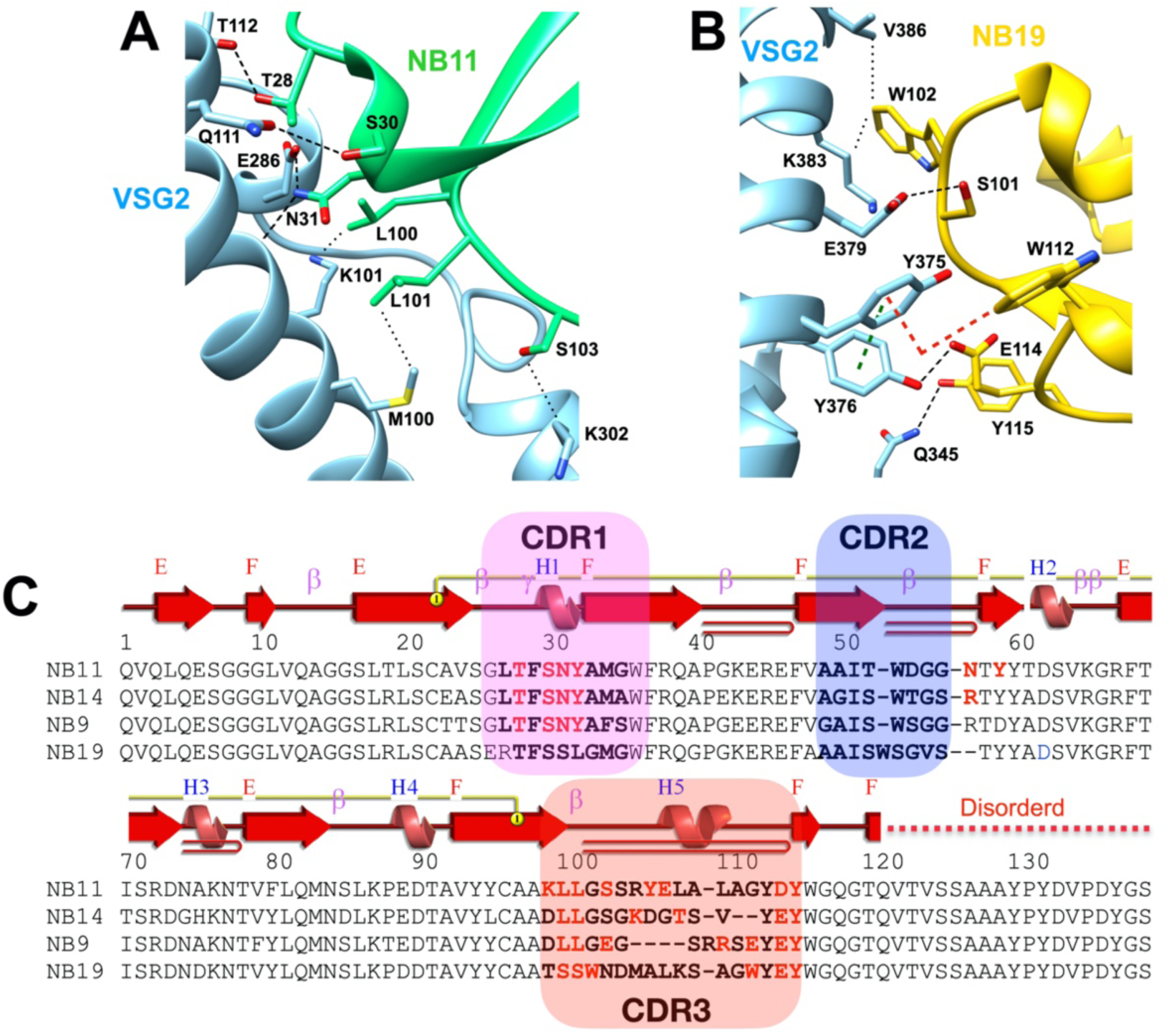
Interactions of Nanobody CDR Variable Loops with VSG2. **(A)** Close up of the interaction of the CDR3 loop from NB11_VSG2_ (NB11) with VSG2. Sidechains from each protein have carbon atoms colored to match the main chain’s ribbon diagram with atoms of oxygen, nitrogen, and sulfur shown in red, blue, and yellow, respectively. Hydrogen bonds (black bricks) are formed between T28, S30 and N31 (NBs) with T112, Q111 and E286 (VSG2), respectively. Residues L100, L101 and S103 of NB11 were found to form non-covalent (black dots) bonds with K101, M100 and K302 of VSG2. **(B**) Close up of the interaction of the CDR3 loop from NB19 with VSG2. Hydrogen bonds (black bricks) are formed between S101 (NB19) and E379 (VSG2), as well as between Y115 (NB19) and Q345 (VSG2). Residues W102 (NB19) and V386 and K383 were involved in non-covalent interactions. W112 of NB19 interacts with Y375 of VSG2 in a parallel displaced π-stacking (red bricks) manner. This interaction is further reinforced by a VSG2 intra-molecule T-shaped π-stacking (green bricks) between Y375 and Y376. **(C)** Sequence alignment and annotation of the four Nbs (numbering is for NB11_VSG2_), including secondary structural elements is shown together with highlighted regions of the CDR variable loops (colored as in Figure 1b). Residues that make contacts with VSG2 are shown with red letters. Structures illustrated with Chimera (Pettersen et al., 2004) and secondary structure with PDBSum (de Beer et al., 2014).

### Nanobody Binding is Detectable on Living Parasites

In order to examine the physiological relevance of these nanobodies and their unexpected binding sites on VSG2, we performed a series of binding experiments to living trypanosomes. Nanobodies harboring a C-terminal hexa-histidine (6xHis) tag were added to trypanosomes expressing different VSGs and the cells analyzed with an anti-tag monoclonal antibody by fluorescence-activated cell sorting (FACS, Methods, Figure 3). Nanobodies NB9_VSG2_, NB11_VSG2_, and NB14_VSG2_ were easily detected as binding to VSG2 expressing cells with several logs difference in signal (Figure 3A), whereas nanobody NB19_VSG2_ could not be detected. In contrast, nanobodies did not bind other VSGs (Figure 3, B-D), thereby showing high specificity (NB11_VSG2_ shown as an example). Verifying that the surface accessibility is not only limited to the relatively small nanobodies, we created a sortaggable variant of the camelid heavy chain IgG2 antibody with the corresponding NB11_VSG2_ Ig domain and found it can, in fact, bind the live parasite coat (Figure S3A).

**Figure 3.**
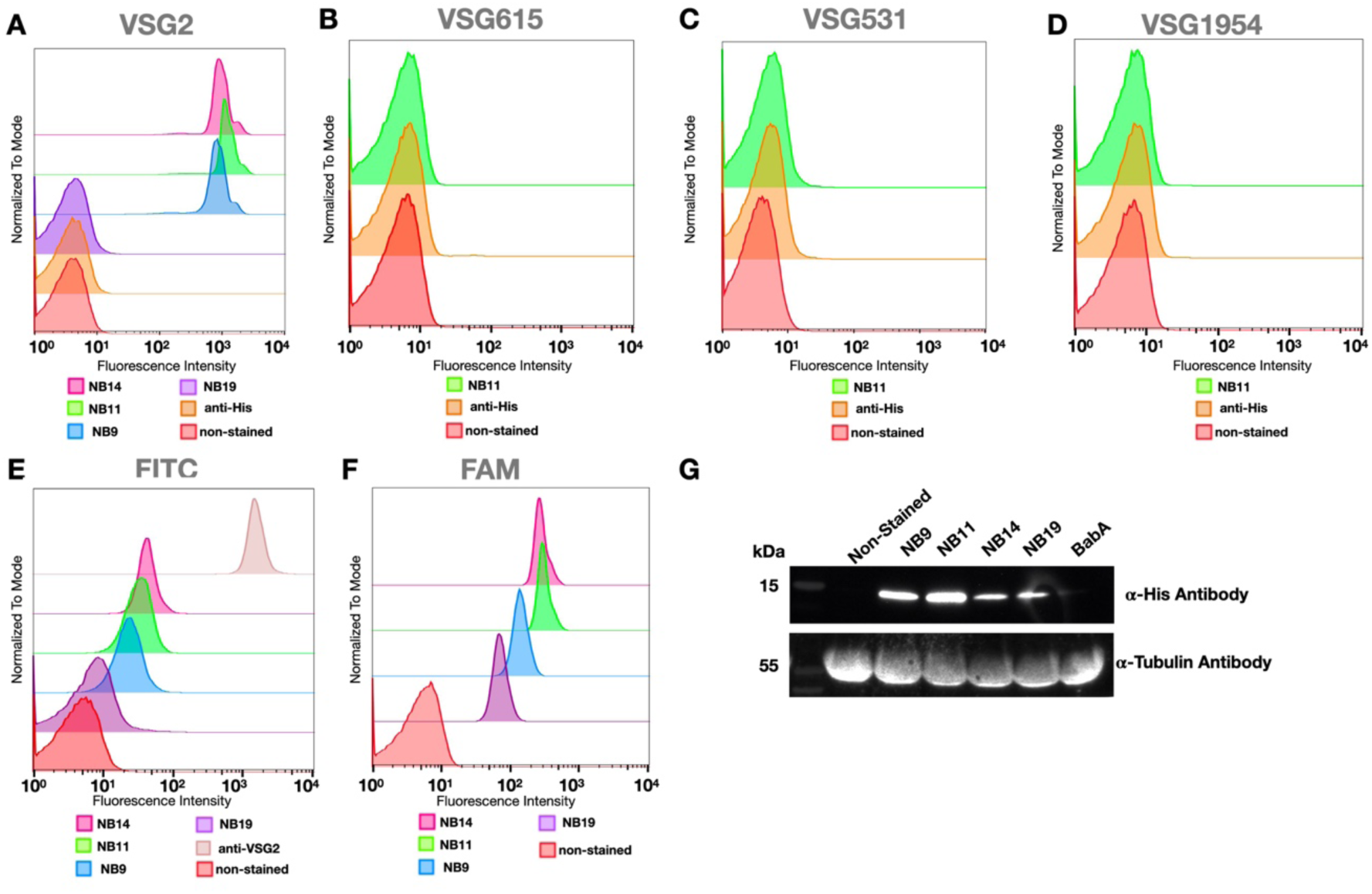
Binding Assays of Nbs to Live Trypanosomes. Different VSG-expressing *T. brucei* were incubated with various Nbs and visualized in different ways by FACS. (A) VSG2-expressing bloodstream form (BSF) trypanosomes were incubated with C-terminally His-tagged Nbs and visualized using an anti-His antibody, (B) VSG615- expressing BSF trypanosomes (expressing a metacyclic VSG) were incubated with C-terminally His-tagged Nbs and visualized using an anti-His antibody, (C) VSG531-expressing BSF trypanosomes were incubated with C-terminally His-tagged Nbs and visualized using an anti-His antibody, (D) VSG1954-expressing BSF trypanosomes (expressing a metacyclic VSG) were incubated with C-terminally His-tagged Nbs and visualized using an anti-His antibody, (E) VSG2-expressing bloodstream form (BSF) trypanosomes were incubated with FITC-labeled Nbs and visualized through the fluorochrome’s emission, (F) VSG2-expressing bloodstream form (BSF) trypanosomes were incubated with Nbs sortagged with FAM and visualized through the fluorochrome’s emission (G) VSG2-expressing bloodstream form (BSF) trypanosomes were incubated with C-terminally His-tagged Nbs, washed thoroughly, run on an SDS-PAGE gel, and probed with an anti-His antibody. An irrelevant Nb (BabA, raised to a protein from *H. pylori*) was used as a negative control. The same cells were probed with an anti-tubulin antibody to provide a gel-loading standard.

The inability to detect NB19_VSG2_ raised the possibility that the binding we observed in solution and in the crystal structure was an artifact of using the soluble VSG2 as the antigen for immunization. A second possibility was that the deep binding we observe in the crystal structure buries NB19_VSG2_ beyond the reach of IgG antibodies. To distinguish between these two models, we directly labeled NB19_VSG2_ and other nanobodies with fluorescent moieties. In one case we chemically conjugated the fluorophore FITC to the proteins (to free amine groups, typically exposed lysine residues), showing that the nanobodies generally could be detected and also how they compare to a full anti-VSG2 IgG (Figure 3E). While a small shift in fluorescence intensity can be observed for NB19_VSG2_-FITC (compared to the non-stained sample, Figure 3F), it showed significance in 2-tailed t-test (p= 0.0109, Figure S3M). To further probe binding, we sought to avoid whole-scale labeling of proteins chemically, and turned to a single, site-specific covalent modification of the Nbs (Figure 3F, Methods). Here, we used the transpeptidase enzyme sortase to attach peptide-linked 6-Carboxyfluorescein (FAM) to the C-terminus of the nanobodies (Methods). In this case, all nanobodies showed binding to the living trypanosomes, although NB19_VSG2_ and NB9_VSG2_ seem to have reduced relative fluorescence intensity (yet with signal intensity over negative controls, Figure 3F). To buttress the positive binding result for NB19_VSG2_ by a third independent experiment, we also exposed living trypanosomes to the nanobodies, thoroughly washed the cells, and then probed them by Western blot (Figure 3G). All the nanobodies, including NB19_VSG2_, were shown to have bound, whereas an irrelevant nanobody raised to the *Helicobacter pylori* BabA antigen did not bind VSG2 expressing trypanosomes.

### Probing Nb-VSG2 Interactions

To further examine the interactions of the Nbs with VSG2, we created mutations in the consensus site of CDR3 of NB11_VSG2_ and two critical contacts to VSG2 from NB19_VSG2_ (Figure 4). We focused on these two because NB11_VSG2_ possess a highly similar consensus binding site to NB9_VSG2_ and NB14_VSG2_, whereas NB19_VSG2_ differs in the binding location and in sequence to the others. The alteration of these important contacts led to a disruption of binding to VSG2 as assayed by gel filtration chromatography. In particular, even single loss-of-contact mutations to the NB19_VSG2_ interface completely disrupted binding, whereas it required a triple mutant in NB11_VSG2_ to abrogate complex formation, suggesting that the complex was more stable (Figure 4).

**Figure 4.**
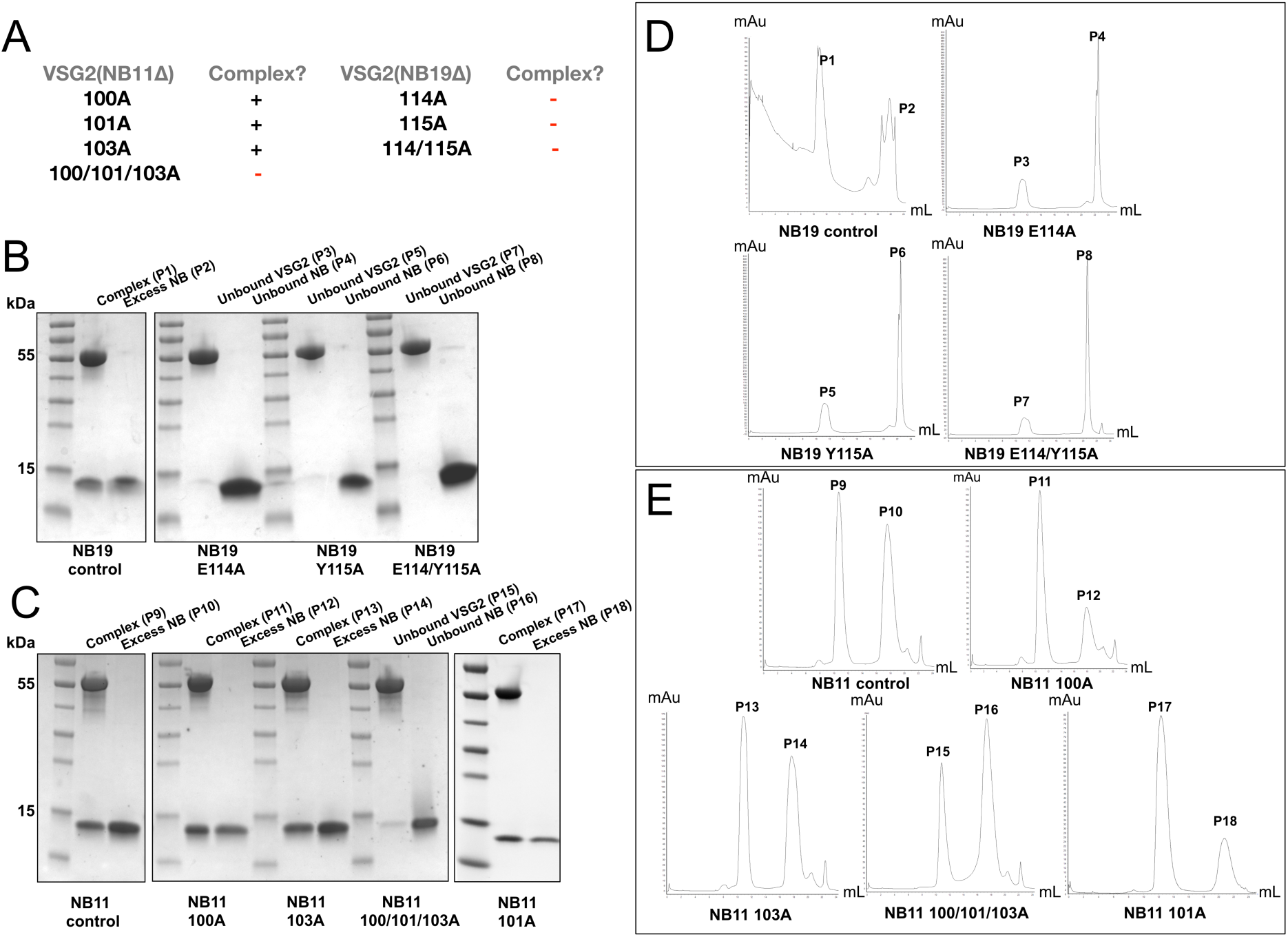
Mutations in the V-Loop Regions of Nanobodies Inhibit Complex Formation with VSG2. **(A)** Overview of NB11 and NB19 mutants in relation to their ability to bind VSG2. NB11 mutants L100A, L101A and S103A form a complex with VSG2 (+), while the triple mutant L100/L101/S103 does not bind (-). For NB19 all single and multiple mutants (E114A, Y115A, E114/Y115A) fail to form a complex with VSG2 (-). **(B and C)** SDS-PAGE analysis of purified, mixed VSG2 and nanobodies (wild type and mutant NB11_VSG2_ and NB19_VSG2_) analyzed by gel filtration chromatography. Peak elution fractions are shown and numbered. VSG2 corresponds to the 55 kDa band and nanobodies to the 15 kDa band. Gels were stained with Coomassie blue. **(D and E)** FPLC chromatographs corresponding to the SDS-PAGE gels **(B and C)** with corresponding peak numbers from the SDS-PAGE gels shown. The Y-axis displays the UV_280nm_ absorbance and the X-axis the elution volume.

### Nanobodies to VSG2 Reduce Coat Mobility, Cause Stiffening of the Cell Body and Stall Parasite Motion

As previous reports had shown Nb-induced interference with trypanosome movement (Stijlemans et al., 2011), VSG2 expressing wild type cells were treated with saturating amounts of the four nanobodies. Only treatment with NB11_VSG2_ resulted in reduced cell motility (Figure S4). While at 5 min post addition of NB11_VSG2_ swimming behavior of the parasites remained largely normal with individual cells actively swimming and changing direction, from 15 min onwards, cell movement was clearly affected as trajectories became shorter and mostly unidirectional (Figure 5A and Figure S4C). A closer inspection of these trajectories suggested that the parasites were not motionless, but that translocation was largely due to drift in the sample which is unavoidable in recordings of this duration. Both swimming speed means and track displacement lengths were extracted from the acquired data (Figures 5B, S5, and S6). Figure 5B (left panel) shows that the mean velocity of NB11_VSG2_ treated cells was reduced compared to that of untreated cells, while NB9_VSG2_, NB14_VSG2_ and NB19_VSG2_ treated cells behaved largely like the control. A comparison of the track displacement length per second also showed smaller values with a narrower range of distribution in NB11_VSG2_ treated cells (Figure 5B, right panel).

**Figure 5.**
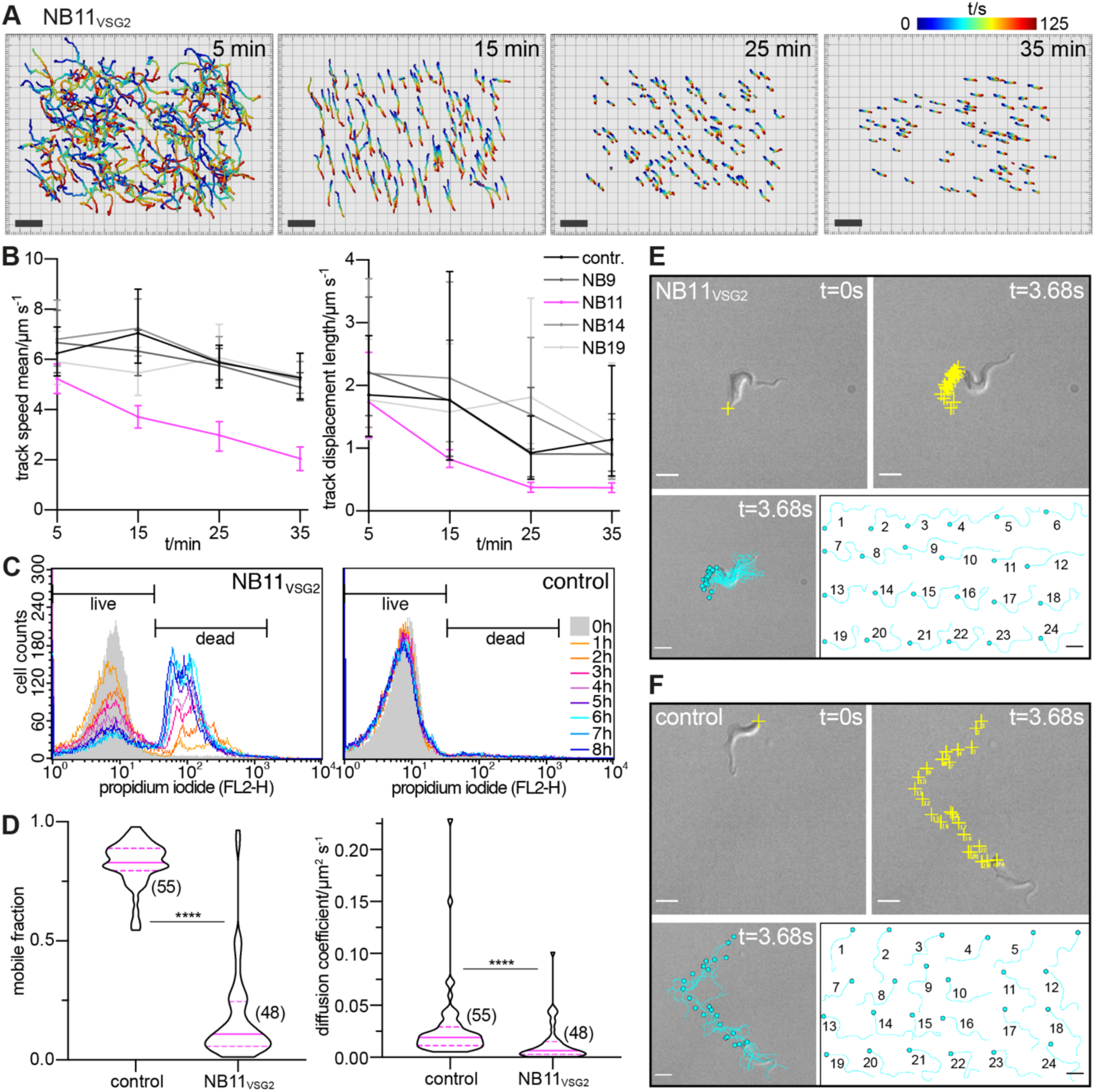
Nanobodies Reduce VSG Mobility, Paralyze, and Kill Trypanosomes. **(A-E)** Cells were treated with a 1.7-fold molar excess of Nb to VSG monomer. **(A)** NB11_VSG2_ affects cell motility after 15 min of treatment. Individual cells in a population were tracked for 125 s at 5, 15, 25 and 35 minutes post addition of NB11_VSG2_ and analyzed using the Imaris x64 software. Cells close to the border of the imaging region were removed by filtering to simplify the images shown (s. Figure S4C for trajectories of complete data set). Trajectories are color coded blue (t = 0 s) to red (t = 125 s). Scale bars, 100 µm. **(B)** Comparison of velocities (track speed mean), left, and track displacement lengths per s, right, of untreated cells and cells treated with four different nanobodies, NB9_VSG2_, NB11_VSG2_, NB14_VSG2_ and NB19_VSG2_. Data extracted from trajectories shown in Figure S4 (median with bars representing the 25th and 75th percentile of data points; see Figures S5 and S6 for scatter dot plots of the data). Both velocity and track displacement are clearly most affected upon treatment with NB11_VSG2_. Flow cytometry analysis to determine the extent of cell death over time following treatment with NB11_VSG2_. Left panel, treated cells, right panel, untreated control cells analyzed in hourly intervals over a duration of 8 h. Propidium iodide was used as a dead cell marker. Each histogram is based on 25.000 gated events (Figure S7 shows the gate settings for the Nb treated data set). The effect of nanobody binding on VSG diffusion was analyzed by fluorescence recovery after photobleaching (FRAP) experiments on immobilized trypanosomes that were surface labeled with ATTO 488. Untreated (control) and NB11_VSG2_ treated cells were analyzed (n in brackets). Violin plots showing the frequency distribution of the mobile fractions, left, and the diffusion coefficient, right, of the VSG in control and NB11_VSG2_ treated cells (median, solid magenta lines; 25th and 75th percentile, dashed magenta lines). There was a remarkable reduction in mobile fraction of treated cells, median of 11 %, compared to that of untreated cells, median of 83 %. The median diffusion coefficient of mobile VSGs following treatment with NB11_VSG2_ was reduced to 0.007 µm^2^/s from 0.019 µm^2^/s for untreated cells. P values from an unpaired, two-tailed Mann-Whitney test (**** = P<0.001) are shown in both graphs. **(E, F)** Single cell analysis of an NB11_VSG2_ treated **(E)** and an untreated **(F)** trypanosome. The upper images show the first (left) and last (right) frame of a video (s. Videos S1 and S2) with the posterior of the cell marked by a yellow cross in the first image and the posterior of every 40th frame of the sequence marked in the second image. Whereas the flagellar beat generates a propulsive force in wt cells the Nb treated cells initiate beats without the generation of a propulsive force. Movement is visualized by tracing the cell shape (cyan lengthwise line through the center of the cell with a circle marking the posterior) in every 40th frame, bottom row. Bottom left image, traces at the position of the cell in the video sequence, bottom right images: the same traces displayed individually to show differences in cell shape and orientation. Scale bars, 5 µm.

As productive movement of trypanosomes was hampered under the influence of NB11_VSG2_, the viability of these cells was analyzed in hourly intervals for 8 hours via flow cytometry (Figures 5C and S7). One hour after NB11_VSG2_ addition at least 80 % of treated cells were still alive, with the proportion of dead cells increasing continuously up to 8 hours when about 25 % of cells were still alive (Figure 5C and Table S3). The amount of nanobody used had a critical impact on cell death (Figure S8, Table S3). A 1:1 molar ratio of NB11_VSG2_ to VSG2 monomer was required and sufficient to induce cell death with a sharp rise in the number of dead cells observed between samples with a 0.9- and 1.1-fold excess, whilst higher Nb concentrations did not alter the kinetics of the cellular response, suggesting that the VSG coat had been saturated with nanobody. Based on these results all further experiments were carried out with a saturating 1.7-fold excess of nanobody to VSG monomer.

We first explored whether nanobody binding might affect VSG diffusion in the cell surface coat. Fluorescence recovery after photobleaching (FRAP) experiments were performed on immobilized cells whose surface proteins had been fluorescently labeled. VSGs of untreated cells exhibited a median mobile fraction of 83 % and diffusion coefficient of 0.020 µm^2^/s. Addition of the Nb resulted in a marked reduction of the median mobile fraction to 11 % with the diffusion coefficient reduced to a median of 0.007 µm^2^/s (Figure 5D, left and right graph, respectively). Hence, binding of NB11_VSG2_ to VSG2 on the cell surface significantly affected the mobility of the protein in the cell surface coat.

As depicted in Figure 5A, a severe restriction in motility was observed in ensemble measurements between 5 and 15 min after addition of NB11_VSG2_. This time period was therefore chosen to acquire high-speed videos of treated and untreated individual cells (Figures 5E and F, Videos S1 and S2). Nanobody-treated parasites were unable to swim productively. The flagellar wave initiated at the tip of the flagellum, but did not progress through the whole cell body so that the typical cellular waveform could not be observed (Bargul et al., 2016). The parasites were partly paralyzed with increased stiffness and decreased ability to bend the cell body. This observation led us to inspect the nanobody treated cells by electron microscopy.

### NB11_VSG2_ Induces the Formation of Nanovesicles and Nanotubes on the Trypanosome Surface

Interestingly, scanning electron microscopy (SEM) of NB11_VSG2_ treated VSG2-expressing cells revealed the formation of nanovesicles and nanotube-like structures as early as 2 minutes following addition of the nanobody (Figure 6, A and B). Initially, nanovesicles of around 200 nm diameter originated mainly from the flagellum attachment zone (FAZ), the flagellar tip (FT) and the posterior pole (PP) of the cell (Figure 6, A and B; Figure S9A, middle and lower inset). In some parasites, the nanovesicles had formed vesicle chains (Figure 6B). In addition to these nanovesicles and vesicle chains, nanotubes of slightly smaller diameter, that could be more than 1 µm in length appeared perpendicular to the FAZ (Figure S9A, upper inset). In the vast majority of cells, nanovesicles and/or nanotubes had been formed after 2 min, while after 5 min all parasites revealed this phenotype. Vesicles appeared on both sides of the FAZ and frequently at the origin of the flagellum, the flagellar pocket (FP) (Figure S9, B and C). Likewise, nanotubes were present in the vicinity of the FP (Figure S9C, left inset). Vesicles in close proximity to the FP tended to be larger (275 nm) than those appearing at the FAZ (150 - 180 nm). In some trypanosomes, the nanovesicles appearing alongside the FAZ seemed to fuse (Figure S9C, right inset), or to redistribute towards the flagellar membrane. However, after 10 min of nanobody incubation, nanovesicles and nanotubes were still present with roughly the same abundance, and, with very little indication of a transition of nanovesicular chains to nanotubes suggesting that both structures form independently (Figure 6C). After 15 min, the number of budding nanovesicles had decreased, and the nanotubes were greatly elongated, thereby apparently forming bridges between parasites. Some of the tubes seemed to be under tension (Figure S9D), while others formed catenated structures.

**Figure 6.**
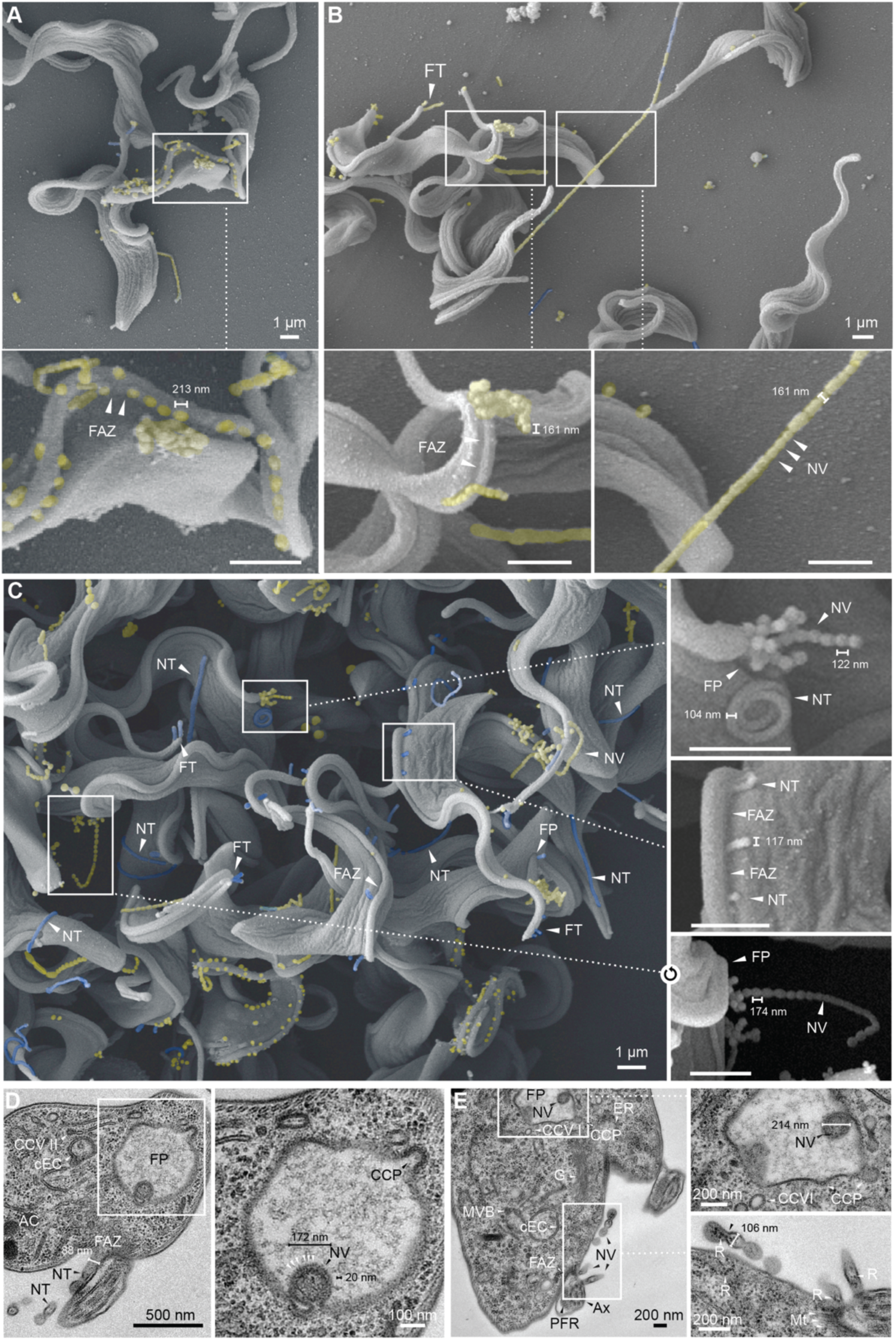
Nanobody Treatment Triggers Formation of Nanovesicles and Nanotubes. **(A-C)** Scanning electron micrographs of trypanosomes treated with a 1.7-fold molar excess of NB11 compared to VSG monomer present. Cells treated for only 2 min **(A,B)** displayed nanovesicles (yellow) as well as some chains of nanovesicles (yellow) with diameters of around 200 nm. FT, flagellar tip; FAZ, flagellar attachment zone; NV, nanovesicle; NT, nanotube; FP, flagellar pocket. Nanotube-like structures (blue) of slightly smaller diameter and lengths exceeding 1 µm could also be observed. All structures emanated from around the FAZ, FT and PP. Cells treated for 10 min **(C)** show an increased and roughly equal abundance of both nanovesicles and nanotubes suggesting them to be independent structures. **(D,E)** TEM images of chemically fixed VSG2 expressing cells treated with a 1.7-fold molar excess of NB11_VSG2_ compared to VSG2 monomer present. CCVII, clathrin coated vesicle II; cEC, circular exocytic carrier; FP, flagellar pocket; AC, acidocalcisome; FAZ, flagellar attachment zone; NT, nanotube; NV, nanovesicle; CCP, clathrin coated pit; ER, endoplasmic reticulum; CCVI, clathrin coated vesicle I; G, Golgi; MVB, multivesicular body; PFR, paraflagellar rod; Ax, axoneme; Mt, mitochondrion; R, ribosome. **(D)** Concomitant formation of a nanovesicle in the flagellar pocket and a nanotube from the proximity of the flagellar attachment zone (FAZ). Formation of a clathrin coated pit in the FP is also apparent and suggests the endocytic pathway to be functional. The budding NV has approx. 20 nm thick structures coating the outer membrane consistent with the length of VSG molecules. **(E)** Concomitant formation of a nanovesicle in the flagellar pocket and nanotubes from the proximity of the FAZ. Nanovesicles can form chains with continuous content and contain electron-dense particles that resemble ribosomes found in the cytoplasm.

Occasionally, smaller nanotubes or nanovesicles appeared to branch off the main tube (Figure S9D, lower inset). The fact that most of the tens of micrometer long tubes were intact, without showing breakpoints suggests that they are mechanically rather robust. This can be seen when tubes bend around trypanosomes (Figure S9D, middle inset). These rigid membrane ropes may well have additionally hampered trypanosome motility, as the parasites can be literally trapped in a nanotubular network. A closer inspection of the apparent network built from nanotubes revealed that most intersections in fact are crossings of two tubes without fusion (Figure S10A, upper inset), while other intersections seemed to represent true membrane fusion (Figure S10A, lower inset). The presence in nanotubes of nodes of similar size as nanovesicles is first seen after 20 minutes (Figure S10A, middle inset). The endpoint of our observation window was 25 minutes, when the nanotubes had formed concatenated or branched networks that are very difficult to trace by SEM (Figure S10B). However, it seems noteworthy that at this time point nanovesicles also emanated from the same trypanosome membrane areas as after 2 minutes of nanobody incubation, namely along the FAZ, at the posterior pole and at the flagellar tip (Figure S10B). Interestingly, when we looked at control parasites that had undergone the same experimental procedure, but without exposure to the nanobody, we occasionally also observed nanovesicle formation at the FAZ, the FT and the FP. Despite most trypanosomes displaying no or few nanovesicles (Figure S10C, 25 min), short nanovesicular chains could be observed in some parasites (Figure S10C, 10 min). This suggests that the formation of nanovesicles at distinct cell surface subcompartments is a biologically relevant process that can be triggered in a spectacular manner by anti-VSG nanobodies (Figure S11).

### Nanovesicles and Nanotubes are Coated with VSGs and Bud from the Cell Surface

To obtain more detailed information about the ultrastructure of the nanovesicles and nanotubes, ultra-thin sections of trypanosomes, treated with nanobody for 15 min, were prepared for transmission electron microscopy (TEM). The micrographs revealed an intracellular architecture that was indistinguishable from that found in non-treated parasites. Figure S12A shows perfect preservation of flagellar pocket structure, including the tight junction at the flagellar pocket collar, the flagellar pocket matrix and the VSG coat. Frequently, clathrin coated pits were visible, indicating that the endocytosis machinery was at least structurally not impaired. Circular and flat endosomal cisternae were present, as well as class 2 clathrin coated vesicles (CCV II), which are essential for cargo transport to late endosomes (Figure 6D) (Grünfelder et al., 2003). Likewise, mitochondrion, nucleus, endoplasmic reticulum, acidocalcisomes, glycosomes, Golgi and the subpellicular microtubule corset did not show any significant alterations (Figures 6E and S12B). Hence, 15 min post-incubation with nanobodies, the parasite’s intracellular architecture was not impaired. Nevertheless, just as in SEM preparations, nanovesicles and nanotubes were readily observable in these TEM samples. These vesicles were covered with an electron-dense layer characteristic of the trypanosome’s VSG surface coat, which was confirmed by immunofluorescence analysis (Figure S13).

Interestingly, nanovesicles were found to be budding into the flagellar pocket. Figure 6D shows a perfectly spherical budding vesicle, that is decorated with evenly spaced structures. The inner diameter of this vesicle is 100 nm, and the outer structures have a size of about 20 nm, which is similar to the size of membrane-embedded VSGs. Note that a clathrin-coated pit is being formed at the same flagellar pocket. Interestingly, the same cell is shedding nanotubes at the flagellar attachment zone. These are smaller in diameter than the FP-derived nanovesicles. From the micrographs, it is not clear how the nanovesicles exit the FP, as the tight junction and flagellar pocket collar are intact (Figure S12B). Figure 6E shows another example of concomitant formation of large nanovesicles within the flagellar pocket and nanovesicles/nanotubes at the flagellar attachment zone. These structures were probably the nanovesicular chains observed in SEM. Interestingly, the TEM images show that the “vesicles” were not always fully separated by membrane but had continuous content (Figure 6E, lower inset). In all these vesicles, electron-dense particles were detected, which were identical with the cytoplasmic ribosomes in size and form (Figure 6E, lower inset). As the flagellum is free of ribosomes, the segmented nanotubes and free nanovesicles likely originate from the pellicular plasma membrane and not from the flagellum (Figure S12D). In contrast, the smooth and long nanotubes observed with SEM after 15 minutes of treatment, could well originate from the flagellar membrane, as shown in Figure S12C. In this cell, multiple tubes bud from one flagellum. These are also covered by a VSG-coat, but do not contain ribosome-like structures. The shedding of the smooth nanotubes must involve significant mechanical forces, as is suggested by marked membrane deformations, both on the flagellar side and on the nanotubes (Figure S12C, white arrows). In rare occasions, a nanotube branchpoint was observed (Figure S12E), but it remains unclear how these branches can form. Figure S12F shows that the tubes can be bent and deformed, which is in agreement with observations made in SEM micrographs.

In conclusion, we have established that nanobodies can bind deep within the VSG-coat and that binding can rapidly affect VSG mobility within the plasma membrane. Interestingly, nanobody-binding leads to an almost instantaneous formation of nanovesicles and nanotubes on the parasite cell surface, concomitant with stiffening of the cells and loss of parasite motility. Cell death occurs significantly later and is probably a consequence of stalled motion, which inevitably will affect cytokinesis.

## Discussion

Using the camelid immune system to generate a collection of anti-VSG2 nanobodies, we have been able to show through structural, biochemical, and cell biological experiments that nanobodies can bind deeply into the trypanosome surface coat, even to the bottom of the NTD of the VSGs. Using secondary IgG antibodies to detect the nanobodies, we also demonstrate that antibodies of the mammalian immune system are able to reach into the densely packed coat of *T. brucei* to a depth of ∼50 Å, but have difficulty reaching further than ∼80 Å below the coat surface (Figure 7A). Furthermore, using full camelid antibodies with the same nanobody domain, we show that the camelid immune system can reach as far as the nanobodies themselves to bind to epitopes buried as deep as 50 Å.

**Figure 7.**
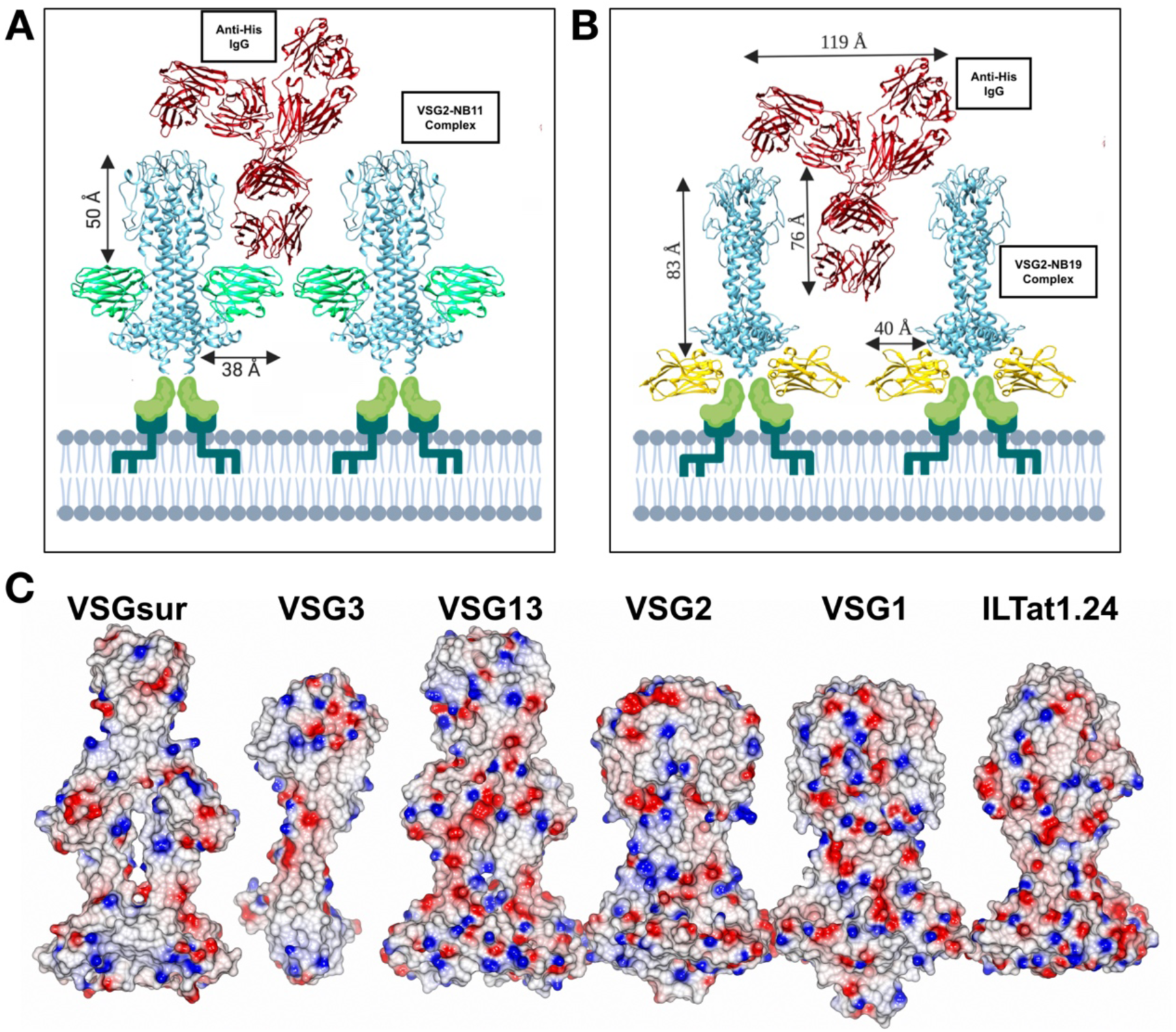
Nanobodies Provide a “Molecular Ruler” to Gauge VSG Accessibility. Schematic of the interaction of an anti-His monoclonal IgG antibody with the VSG coat during FACS experiments. The trypanosome membrane is shown at the bottom with idealized lipids into which the (dark green) GPI anchor of the C-terminal domain (medium green) inserts. Above that is a ribbon diagram drawing of a VSG2 dimer (cyan) with a bound NB (NB9, NB11, NB14) superimposed in light green, **A**) or NB19 (gold, **B**). The IgG is shown as a ribbon diagram colored red (PDB ID 5DK3). **(C)** Panel of crystal structures of VSGs drawn as molecular surfaces colored by relative electrostatic potential (blue is positive/basic, red is negative/acidic, and white is neutral). RCSB PDB IDs for models used are ILTat1.24 (2VSG), VSG1 (5LY9), VSG2 (1VSG), VSG13 (6Z8H), VSG3 (6ELC), and VSGsur (6Z7A). Molecular graphics software used: Chimera (Pettersen et al., 2004), CCP4mg (McNicholas et al., 2011), and biorender.com Pro.

Our experimental results demonstrate that the antigenic surface of the VSGs extends well beyond the upper surfaces of these molecules. Such conclusions are consistent with the observations that the VSGs possess molecular surfaces with relatively high sequence divergence as well as markedly distinguishing surface properties not only at the “top” of the NTD, but across the entire molecule (Figure 7B). That the much more highly conserved CTD would be difficult for antibodies to access (located more than 100 Å below the surface) as shown previously (Schwede et al., 2011) and in our study, is also in harmony with this model.

We identified one Nb (NB11_VSG2_) that was highly efficient in immobilizing and ultimately killing cells, a marked motility phenotype which is in accordance with published work (Stijlemans et al., 2011). In contrast, our work does not support the conclusion that this is due to impaired endocytosis and hence impaired energy supply. The onset of the motility phenotype was within minutes of nanobody addition and electron micrographs did not reveal any disruption in the endomembrane system, including the endosomes. What we observed was a marked reduction in cell body flexibility and increased stiffness of parts of the cell. This was not caused by paralysis of the flagellum as it still initiated the beat which was however not propagated over the cell body. Hence, the trypanosomes did not swim anymore. We hypothesize that the phenotype observed could be related to the mobility and integrity of the surface coat. We have previously shown that the VSG coat operates very close to a molecular crowding threshold, where a high lateral mobility is balanced with the highest possible density of the protein (Hartel et al., 2016). When the VSG density exceeds this threshold, translational diffusion in the surface coat is severely affected. This is exactly what might occur in the experiments reported here. Binding of the nanobody increases the hydrodynamic radius of the VSG dimers which results in a dramatic decrease in the mobile fraction. Thus, rotational and translational diffusion of the VSGs is confined by steric hindrance. Thermodynamics in confinement would cause tension within the VSG coat due to pressure forces. If this were the case, there should be ultrastructural changes in surface topology of the trypanosomes. Indeed, we observed dramatic changes of the trypanosome cell surface in electron micrographs.

Membrane fission is essential for the life of all cells. Cytokinesis, the generation of organelles such as the mitochondrion, ER and Golgi, as well as endomembrane trafficking are processes that all depend on this type of membrane remodeling. Complex machineries involving many proteins that can form helical scaffolds or constricting rings, as well as shallow membrane insertions, are thought to control membrane fission (Snead et al., 2017; Campelo and Malhotra, 2012). Classic examples are the clathrin or COP pathways and the ESCRT machinery. These pathways are well-controlled in space and time, and they are highly energy-consuming. The total energy of membrane vesiculation has been estimated to be in the range of several hundred k_B_T, which means that membrane fission requires significant mechanical forces, acting from outside onto the membrane. This is exemplified by the massive clathrin assembly visible on a coated pit at the flagellar pocket in Figure S12A.

The nanovesicles and nanotubes observed in this work on the trypanosome cell surface did not show any structural features that differed from the plasma membrane. They were decorated with an electron dense VSG layer but no additional coats that could be involved in vesicle formation were detectable. This is not unexpected, as it is very unlikely that the parasites can sense the nanobody addition, transduce this information and actively secrete a putative extracellular fission-promoting machinery - all within seconds. Thus, it is most probably not an active cellular process that drives nanovesicle and nanotube formation.

An attractive, alternative explanation for the phenotype would be protein-protein crowding. The addition of nanobodies to the extremely dense VSG coat could - within seconds - raise the protein density on the cell surface above the molecular crowding threshold. In fact, we have shown that the lateral mobility of the VSG decreases rapidly after nanobody addition, which supports this assumption. With the binding of nanobodies, mobility of the VSG becomes restricted and the coat is therefore under tension. Collisions of nanobody-loaded VSGs will generate pressure (on one side of the plasma membrane). This behavior of a crowded membrane protein has been likened to that of compressed gas and it has been suggested to provide the pressure force required to initiate and complete membrane vesiculation and tubulation (Snead et al., 2017). Thus, nanobody-mediated protein crowding could provide the energy required for membrane fission. No cellular machineries would be required to overcome the energy barrier. Membrane fission by protein-protein crowding has been observed before, albeit only in artificial systems and is supported by modeling approaches (Derganc and Čopič, 2016; Snead et al., 2017).

Our data suggest that this mechanism could also function in the cellular context. Due to its uniformity, the trypanosome coat with its high density and essential mobility of the VSG is an ideal model system to study this new mechanism of membrane fission in living cells. The fact that we find nanotubes and nanovesicles also in untreated cells, albeit only occasionally, suggests that, at least in trypanosomes, protein crowding-mediated extracellular vesiculation could be a biologically relevant process. We surmise that by using this passive, density-controlled process, the parasites can adjust the VSG-coat density, e.g. during cell division or VSG switching. Protein crowding-mediated membrane shedding may well be effective on other cells with dense protein coats.

Especially at early time points after addition of nanobodies, vesicle chains of increasing length were observed. TEM images showed that the chains were not always composed of separated vesicles but were often segmented, continuous volumes. There are few reports on vesicle chains in biology. Outer membrane vesicle chains have been reported in *Myxococcus xanthus* (Remis et al., 2014) and biopearling has been observed in a marine flavobacterium (Fischer et al., 2019). However, the mechanisms underlying vesicle chain formation have not been studied. As segmented membrane volumes are detectable already after 2 minutes in nanobody-treated trypanosomes, we surmise that counteracting mechanical forces interrupt full fission of nanovesicles. The segmented vesicle chains mainly appear at the FAZ and the flagellar pocket. These regions continuously experience the forces exerted by the flagellar beat. It is tempting to speculate that the flagellar force causes interruption of fission events. The protein crowding pressure on the nascent vesicle could be strong enough to maintain the vesicle curvature. When a new vesicle forms at the same position of the FAZ it would be connected to the previous one. In this way, the continuous beat of the flagellum would produce segments of very similar size. After about 15 minutes of nanobody treatment, the vesicle chains disappear. This is the time point when the propagation of the flagellar beat is impaired, probably due to membrane tension caused by loss of membrane through vesiculation. Thus, the force producing the segmented membrane volumes decreases, resulting in a shift to tubulation. The resulting nanotubes can be tens of micrometers long, and the wiggling cells entangle themselves in these rope-like structures. It is important to note that fusion of nanotubes to neighboring parasites has not been observed, suggesting that cell-cell communication via membrane bridges is not occurring.

In summary, the fast onset of membrane fission at “free” membrane compartments could be explained by membrane tubulation through protein-protein crowding on the trypanosome cell surface. Are the nanovesicles in fact exosomes or ectosomes (extracellular vesicles, EVs)? Exosomes are produced by multivesicular bodies and secreted by exocytosis. This has previously been observed in fly-stage *T. brucei* procyclic forms (Eliaz et al., 2017). Since we did not find any changes in the endomembrane architecture, the vesicles formed after nanobody treatment are likely not exosomes. The formation of extracellular vesicles in trypanosomes was first reported in 2016 (Szempruch et al., 2016). In contrast to our data, this work revealed that EVs were generated by vesiculation of nanotubes that are highly fusogenic and originate from the flagellar membrane. Thus, despite morphological similarities, the nanovesicles and nanotubes observed in the present study are neither exosomes nor ectosomes, but rather the product of membrane fission through protein-protein crowding. The trypanosome-nanobody system for the first time enables us to systematically study a highly efficient mechanism by which the crowded protein environment on the surface of cells causes changes in membrane shape.

## SUPPLEMENTARY FIGURES AND TABLES

Figures 1-15, Table 1-3

## ACKNOWLEDGEMENTS

We acknowledge synchrotron time at the Diamond Light Source (DLS, beamline i03, Neil Paterson and colleagues and the Paul Scherrer Institut, Villingen, Switzerland (SLS, beamline PXIII, Vincent Olieric and colleagues). We thank Steve Schoonooghe, Ema Estevens Romão, and Gholamreza Hassanzadeh Ghassabeh of the Vlaams Instituut voor Biotechnologie (VIB) Nanobody Core facility for working closely with us to produce anti-VSG2 VHH antibodies and donating for experimental controls the antibody to *H. pylori* BabA. We thank Elisabeth Meyer-Natus (Würzburg) for technical assistance with EM sample preparation and data acquisition. We thank Tim Krüger (Würzburg) for assistance with microscopy and Susanne Fenz (Würzburg) for helpful discussions. M.E. is supported by DFG grants EN305, DFG-SPP1726, DFG-GRK2157 and GIF grant I-473-416.13/2018.

## AUTHOR CONTRIBUTIONS

C.E.S., F.N.P., M.E. and N.G.J. conceived, initiated and coordinated the project, M.vS. produced VSG2 for production of nanobodies. M.vS. and A.H. produced and crystallized NB11_VSG2_ and NB14_VSG2_ complexes. A.H. produced and crystallized NB9_VSG2_ and NB19_VSG2_ complexes, all complexes for membrane studies (with M.vS. also producing NB11_VSG2_ and NB14_VSG2_ for the latter), collected the diffraction data and solved all structures. J.Z. aided in crystal preparation and beamline data collection. H.H. and A.H. created sortaggable Nbs and the camelid heavy chain Ig2. H.H., M.vS, and A.H performed anti-His FACs binding experiments. H.H and A.H produced the FAM and FITC FACS data on live *T. brucei*, while A.H. executed FACS experiments on non- VSG2 coats. L.H. performed all electron microscopy, parasite movement, coat diffusion, and toxicity experiments. C.E.S., A.H., N.G.J. and M.E. wrote the manuscript. All authors discussed the data and approved the final manuscript.

