## Supplementary Material for "Nanobody Mediated Macromolecular Crowding Induces Membrane Fission and Remodeling in the African Trypanosome"

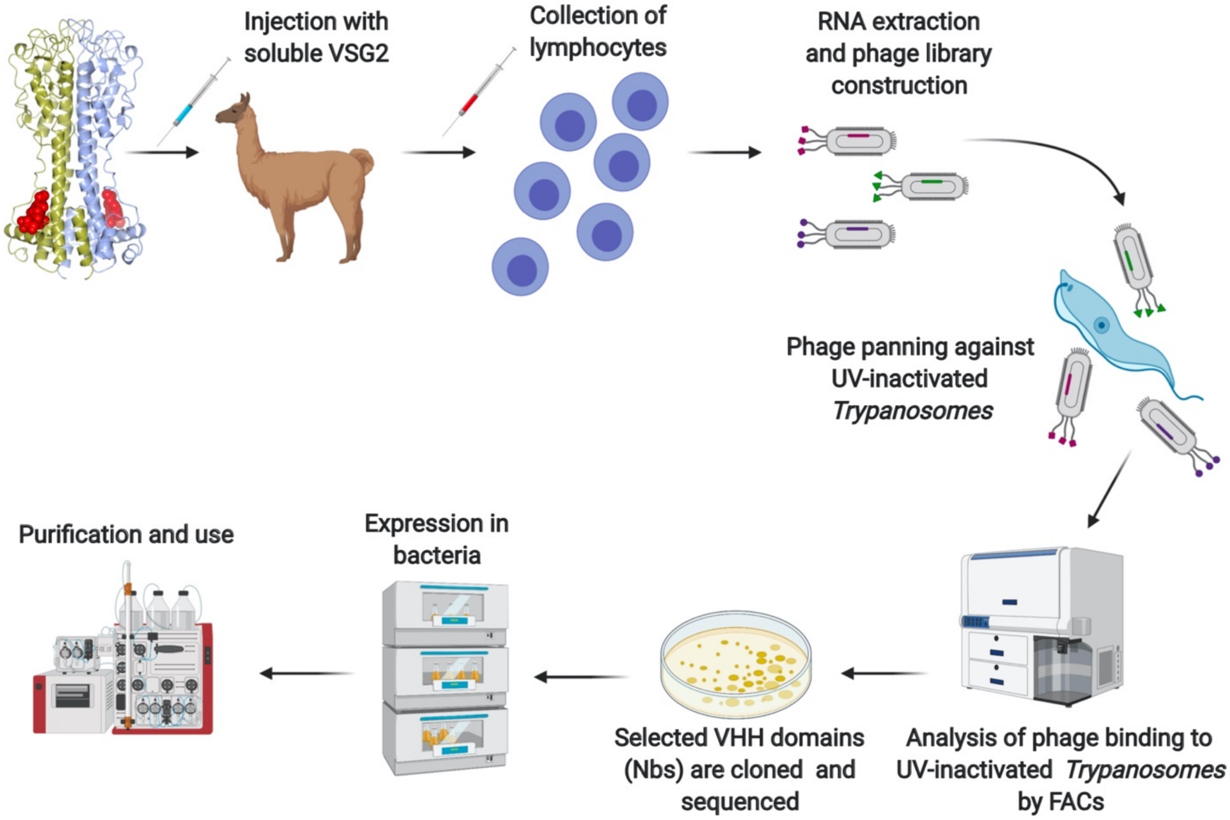

**Figure S1. Production of anti-VSG2 Nanobodies**

Schematic illustrating the broad steps of Nb production (see Methods for details). Figure created with biorender.com Pro.

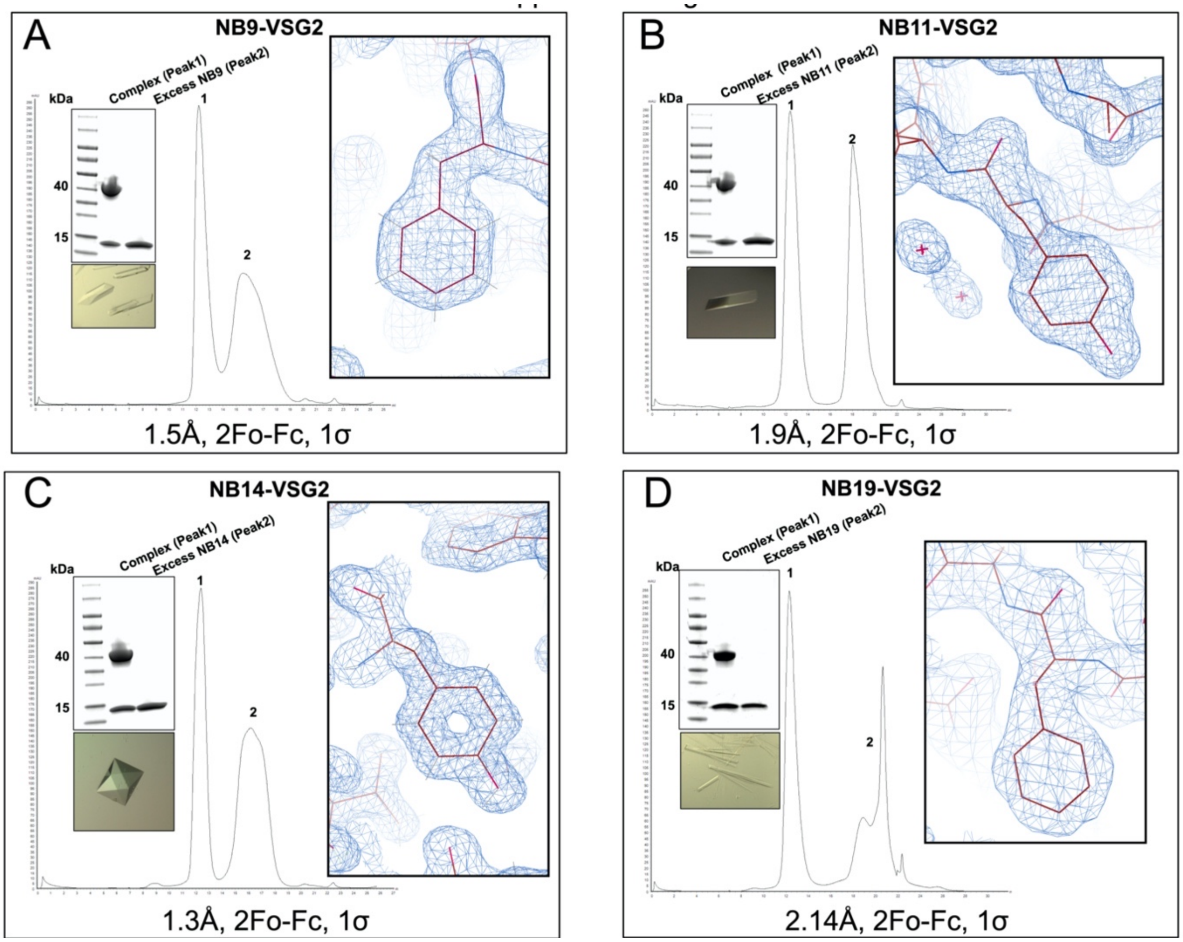

**Figure S2. Purification, Crystallization, and Representative Electron Density of Nb-VSG2 Complexes, Related to Figure 1**

Summary of various steps in the crystallographic structural solution of VSG2 in complex with A) NB9, B) NB11, C) NB14, and D) NB19. Panels showing the gel filtration chromatogram of purified NB-VSG2 NTD and a Coomassie stained SDS-PAGE gel of the final material used for crystallization. Images of crystals grown in hanging drops and final model 2Fo-Fc electron density contoured at 1σ.

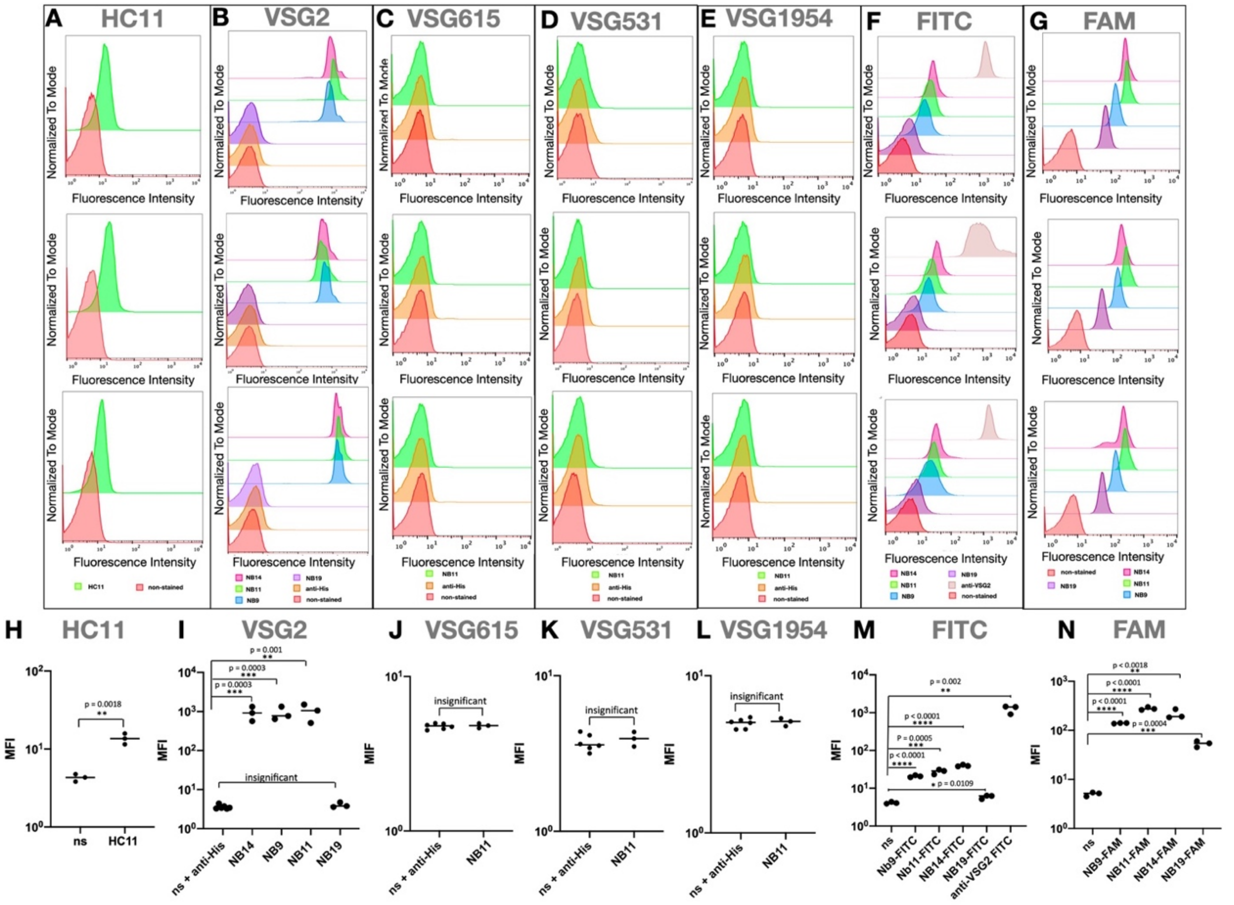

**Figure S3. Triplicates of Different VSG-expressing *T. brucei* Incubated with Various Nbs and Visualized in Different Ways by FACS and Analyzed by Two-tailed T-test.**

(A) The VHH domain from NB11$_{VSG2}$ was reconstructed into full-length Camelid IgG2 heavy-chain (HC) antibodies. 2T1 cells expressing VSG2 were stained (see methods) with FAM-labelled antibodies. Similar to the smaller Nb experiments on live parasites also the HC antibody containing the NB11 domain was able to bind the surface coat. (B) Triplicates of VSG2-expressing bloodstream form (BSF) trypanosomes were incubated with C-terminally His-tagged Nbs and visualized using an anti-His antibody. (C) Triplicates of VSG615-expressing BSF trypanosomes were incubated with C-terminally His-tagged Nbs and visualized using an anti-His antibody. (D) triplicates of VSG531-expressing BSF trypanosomes (expressing a metacyclic VSG) were incubated with C-terminally His-tagged Nbs and visualized using an anti-His antibody. (E) VSG1954-expressing BSF trypanosomes (expressing a metacyclic VSG) were incubated with C-terminally His-tagged Nbs and visualized using an anti-His antibody. (F) VSG2-expressing bloodstream form (BSF) trypanosomes were incubated with FITC-labeled Nbs and visualized through the fluorochrome's emission. (G) VSG2-expressing bloodstream form (BSF) trypanosomes were incubated with Nbs sortagged with FAM and visualized through the fluorochrome's emission. (H) Two-tailed t-test showing a significant difference between the ns fluorescence intensity (FI) (negative controls) and the HC signal (p=0.0018). (I) Two-tailed t-test showing a significant difference between the ns+anti-his (negative control) and the NB14 (p=0.0003), NB9 (p=0.001), NB11 (p=0.001) FI. The difference in signal for NB19 was not significant. (J) Two-tailed t-test showing no significant difference between the ns FI (negative control) and NB11 signal on VSG615 expressing parasites, (K) Two-tailed t-test showing no significant difference between the ns FI (negative control) and NB11 signal on VSG531 expressing parasites. (L) Two-tailed t-test showing no significant difference between the ns FI (negative control) and NB11 signal on VSG1954 expressing parasites, (M) Two-tailed t-test showing a significant difference between the ns (negative control) and the NB9-FITC (p=0.0001), NB11-FITC (p=0.0005), NB11 (p=0.0001), NB19-FITC (p=0.0109) and the anti-VSG2-FITC FI (p=0.002). (N) Two-tailed t-test showing a significant difference between the ns (negative control) and the NB9-FAM (p=0.0001), NB11-FITC (p=0.0001), NB11 (p=0.0018) and NB19-FITC (p=0.0004) FI. FACS data was analyzed and plotted (FlowJo™ Software Version 10.5.3. Ashland, OR: Becton, Dickinson and Company; 2019) and statistical analysis was performed using Prism 8 (two-tailed t-test).

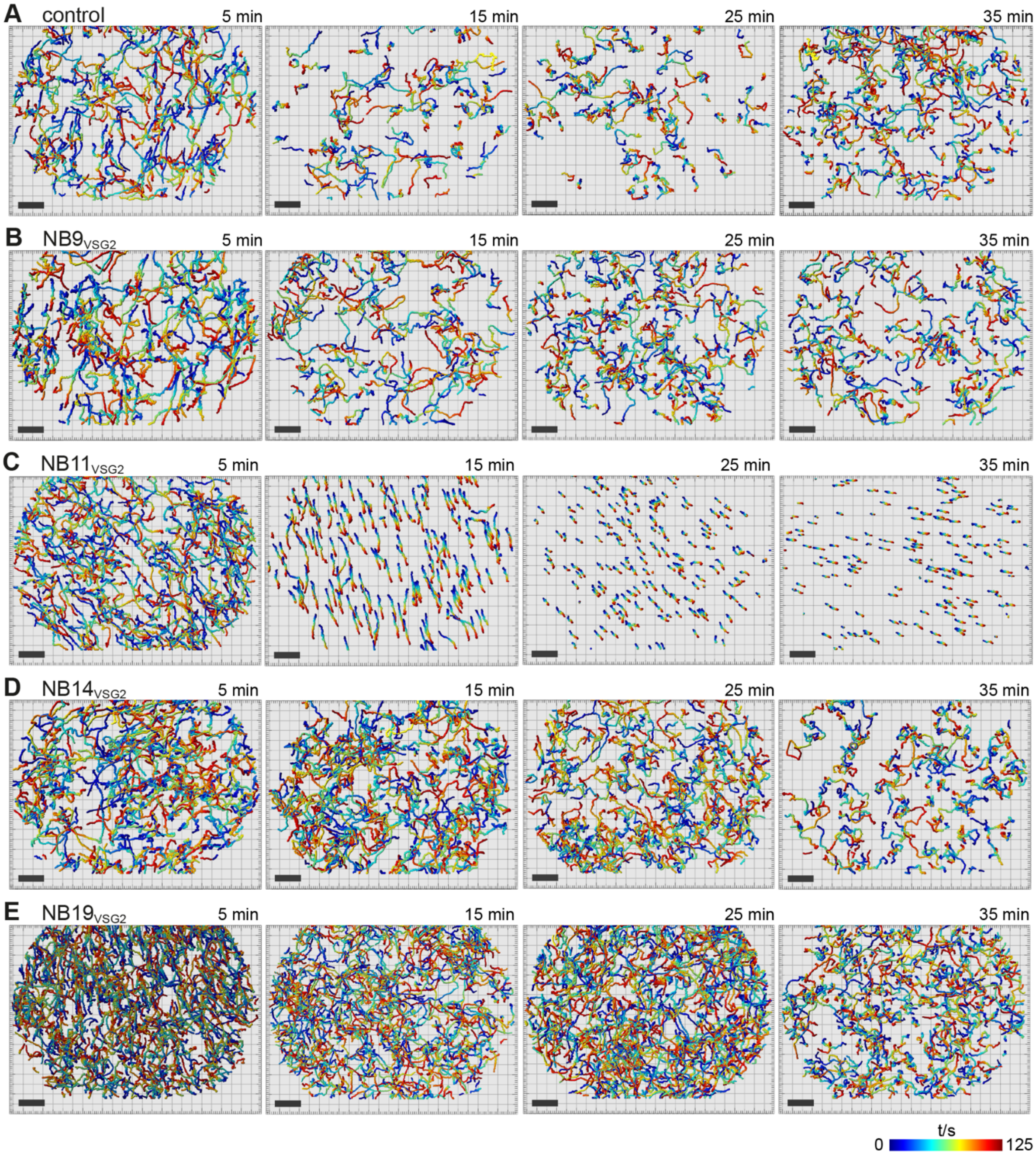

**Figure S4. Trypanosome Motility Following Treatment with the anti-VSG2 Nanobodies NB9, NB11, NB14 and NB19, Related to Figure 5**

(A) Untreated control cells, (B-E) cells treated with a 1.7-fold molar excess of nanobody, (B) $NB9_{VSG2}$, (C) $NB11_{VSG2}$, (D) $NB14_{VSG2}$, (E) $NB19_{VSG2}$. Methylcellulose was added to the cells prior to imaging in order to achieve a viscosity of 25 mPa and ensure ideal swimming conditions for trypanosomes. Images were acquired for a period of 125 s at 5, 15, 25 and 35 min post addition of the respective NB and in the case of the untreated control at the same time points following sample preparation. Trajectories were visualized and analyzed using the Imaris x64 software and are color coded according to the acquisition time (0 s in blue and 125 s in red). Scale bars, 100 μm.

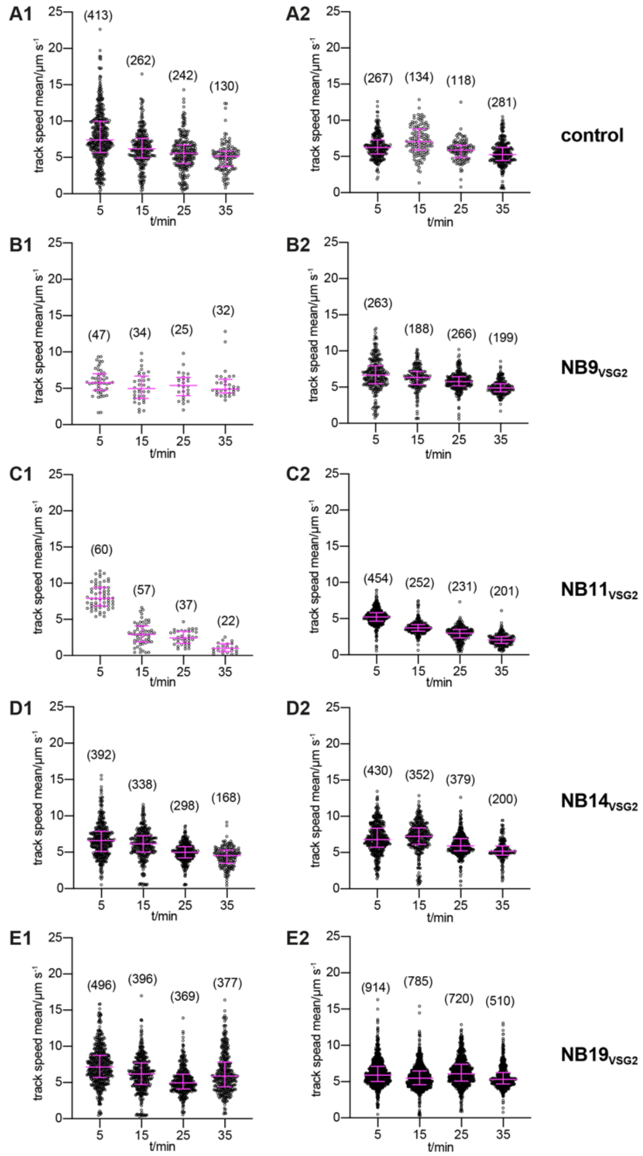

**Figure S5. The Velocity of Trypanosomes is Reduced Following Treatment with Nanobody NB11$_{VSG2}$, Related to Figure 5**

Scatter dot plots showing the track speed mean for trajectories (s. Figure S4) of individual cells at 5, 15, 25 and 35 min time points of untreated control cells (A) and post addition of a 1.7-fold molar excess of anti-VSG2 nanobodies NB9$_{VSG2}$ (B), NB11$_{VSG2}$ (C), NB14$_{VSG2}$ (D) and NB19$_{VSG2}$ (E) compared to VSG monomer present on the cells. Number of trajectories analyzed shown in brackets. The median and 25th and 75th percentile are shown in magenta. The viscosity of samples used for data acquisition was adjusted to 25 mPa with methylcellulose. Data presented in A1-E1 are from a pilot experiment with continuous data acquisition for 30 min, whereas for the data presented in A2-E2, videos were recorded for 125 s specifically at the time points given. Trajectories were analyzed using the Imaris x64 software.

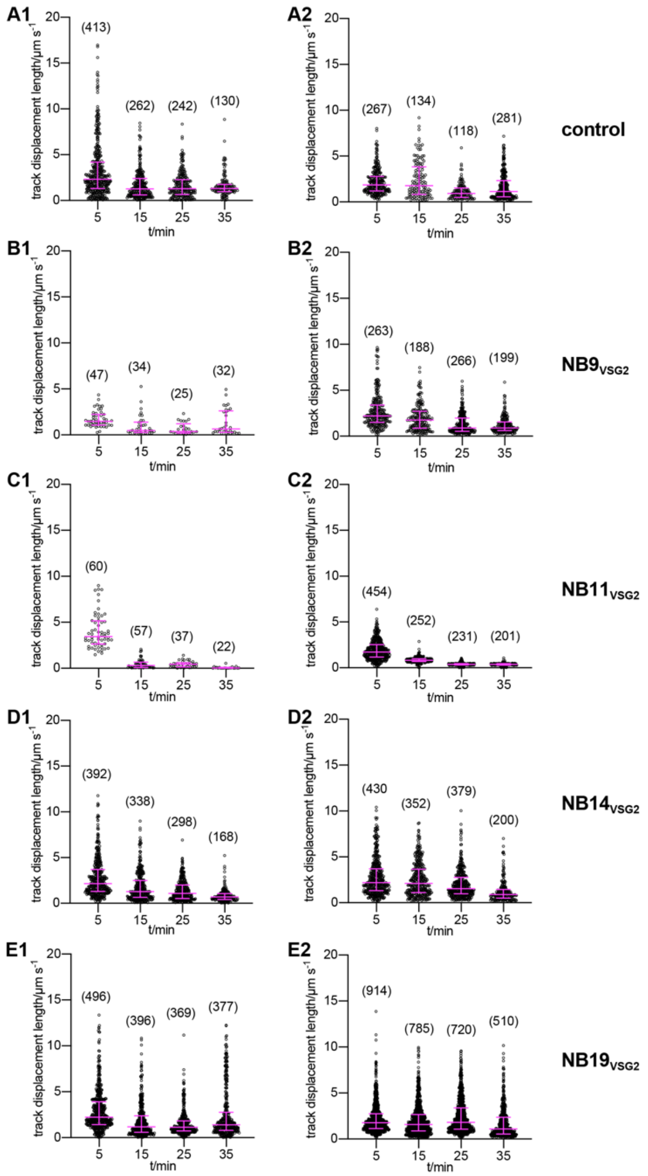

**Figure S6. The Average Track Displacement Length and Distribution of Values within the Population of Trypanosomes is Reduced Following Treatment with Nanobody NB11$_{VSG2}$, Related to Figure 5**

Scatter dot plots showing the average track displacement length per s for tracks of individual cells at 5, 15, 25 and 35 min time points of untreated control cells (A) and post addition of a 1.7-fold molar excess of anti-VSG2 nanobodies NB9$_{VSG2}$ (B), NB11$_{VSG2}$ (C), NB14$_{VSG2}$ (D) and NB19$_{VSG2}$ (E) compared to VSG monomer present on the cells. The median and 25th and 75th percentile are shown in magenta. The viscosity of samples used for data acquisition was adjusted to 25 mPa with methylcellulose. Data presented in A1-E1 are from a pilot experiment with continuous data acquisition for 30 min, whereas for the data presented in A2-E2 videos were recorded for 125 s specifically at the time points given. Tracks were analyzed using the Imaris x64 software.

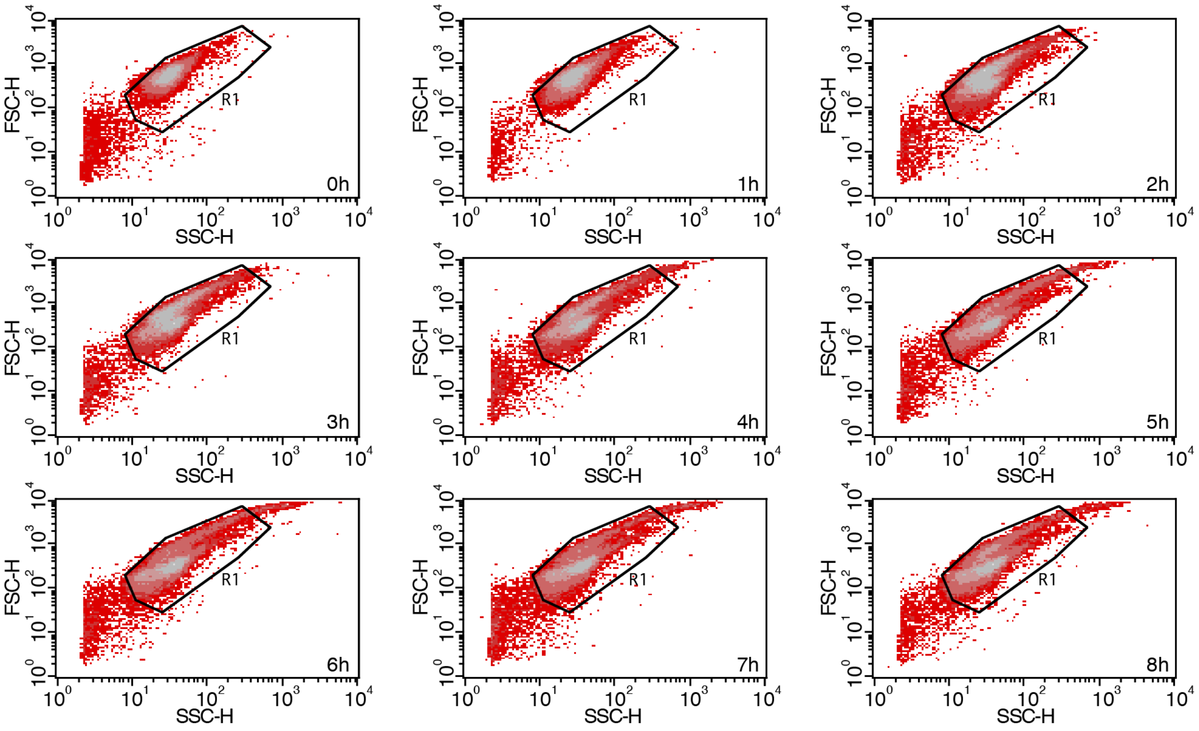

**Figure S7. Scatter Plots from Flow Cytometry Analyses of NB11$_{VSG2}$ Treated Trypanosomes, Related to Figure 5**

Cells were treated with a 1.7-fold molar excess of NB11$_{VSG2}$ compared to VSG monomer present and analyzed hourly between 0 and 8 h for live vs dead cells following treatment with the dead cell marker propidium iodide. Shown are the scatter plots at each time point and the gate R1 used for data acquisition and analysis. 25.000 gated events were recorded for each time point.

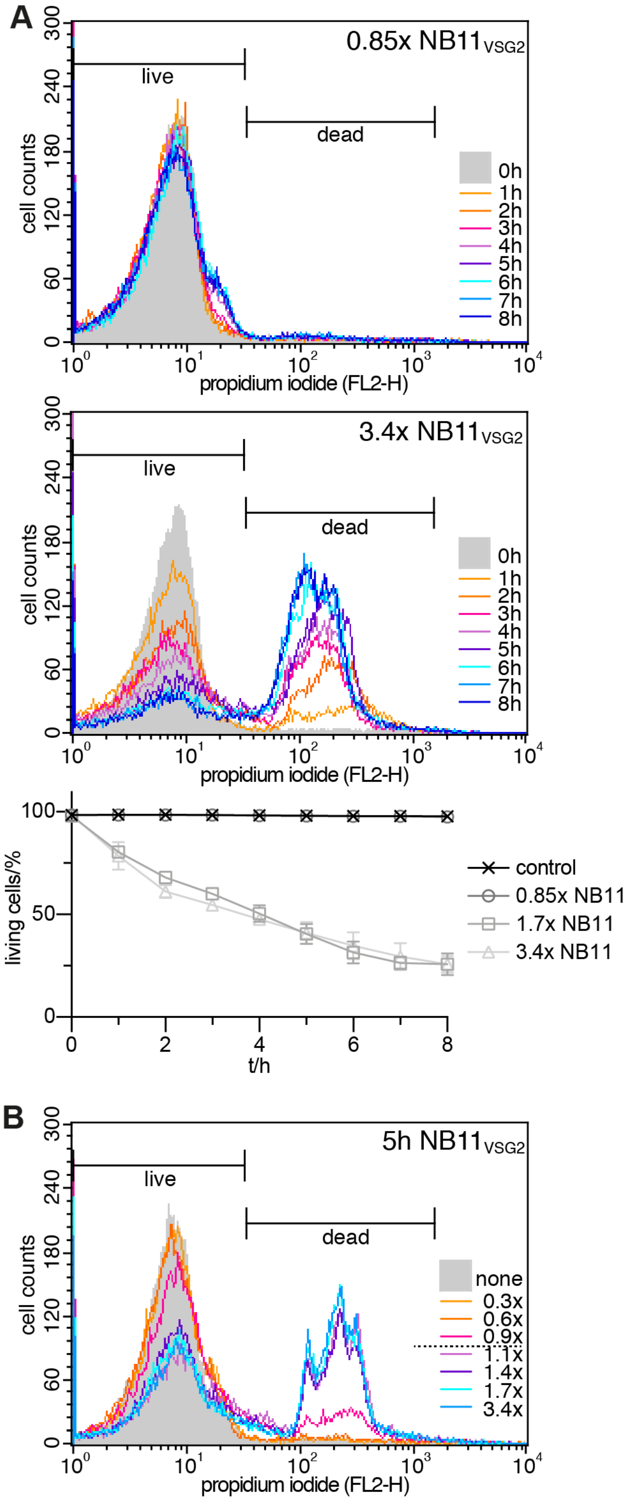

**Figure S8: Cell Death of Trypanosomes Treated with NB11$_{VSG2}$ is Concentration Dependent and Saturates at an Equimolar Amount of Nanobody to VSG Monomer, Related to Figure 5**

(A) Histogram plots of flow cytometry analyses to determine the extent of trypanosome cell death over time following treatment with NB11$_{VSG2}$. In addition to treatment with a 1.7-fold molar excess of nanobody (s. Figure 5C) cells were treated with a 0.85 (top) and 3.4-fold (middle) molar excess. Live and dead cells were distinguished by treatment with the dead cell fluorescence marker propidium iodide (PI) just prior to flow cytometry analyses. 25.000 gated events were recorded at each time point. Whereas cells treated with less than an equimolar amount of nanobody to VSG monomer survived the duration of 8 h (comparable to untreated cells), cells treated with a 3.4-fold molar excess displayed similar kinetics of cell death to cells treated with a 1.7-fold molar excess of NB11$_{VSG2}$. Graph (bottom) displaying the amount of living cells at each time point for untreated cells and cells treated with a 0.85, 1.7 and 3.4-fold excess of NB11$_{VSG2}$. Experiments were carried out in triplicate and data are represented as mean ± SD. Where error bars are not shown they are smaller than the symbol for the data point.

(B) Histogram plots of flow cytometry analyses of VSG2 expressing cells treated with a range of a 0.3 to 3.4-fold molar excess of NB11$_{VSG2}$ to VSG2 monomer over a fixed time of 5 h. Dead cells were distinguished from living cells by the addition of the dead cell fluorescence marker propidium iodide (PI) prior to data acquisition. 25.000 gated events were acquired for each concentration analyzed. The maximum amount of cell death is reached at a molar excess of NB11$_{VSG2}$ to VSG2 monomers between 0.9 and 1.1.

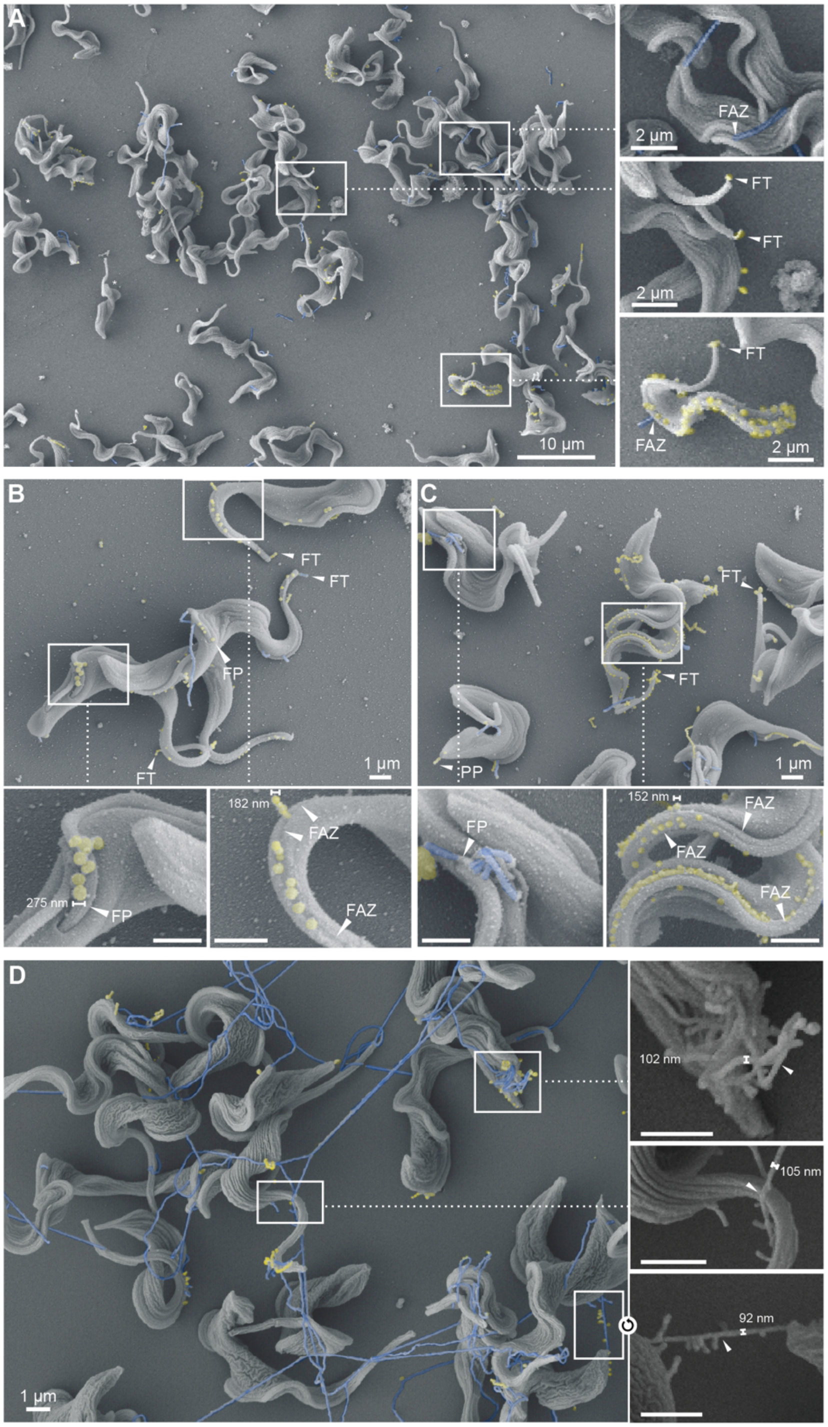

**Figure S9. As Time Progresses NB11VSG2 Leads to Increased Nanotube Formation in VSG2 Expressing Cells, Related to Figure 6** (A-D) SEM images of NB11$_{VSG2}$ treated VSG2 expressing cells. FAZ, flagellar attachment zone; FT, flagellar tip; FP, flagellar pocket; PP, posterior pole.

(A) At 2 minutes post addition of $NB11_{VSG2}$ nanovesicles (highlighted in yellow) are seen originating from the flagellar tip and the vicinity of the FAZ (inset middle and bottom). Nanotubes (highlighted in blue) can also be observed extending from the FAZ (inset top and bottom) and can be several µm in length. (B,C) At 5 min post addition of $NB11_{VSG2}$ both nanovesicles and nanotubes can be observed emanating from the FT, FAZ and the PP of the *T. brucei* cells. Nanovesicle originating from the FAZ (right insets of B and C) are smaller in diameter than those found close to the flagellar pocket (left inset of B). (D) At 15 min post addition of $NB11_{VSG2}$ long nanotubes (blue) extending from the trypanosomes are prominent, though nanovesicles (yellow) are still being formed. The diameter of these tubes lies in the range of 100 nm (see insets).

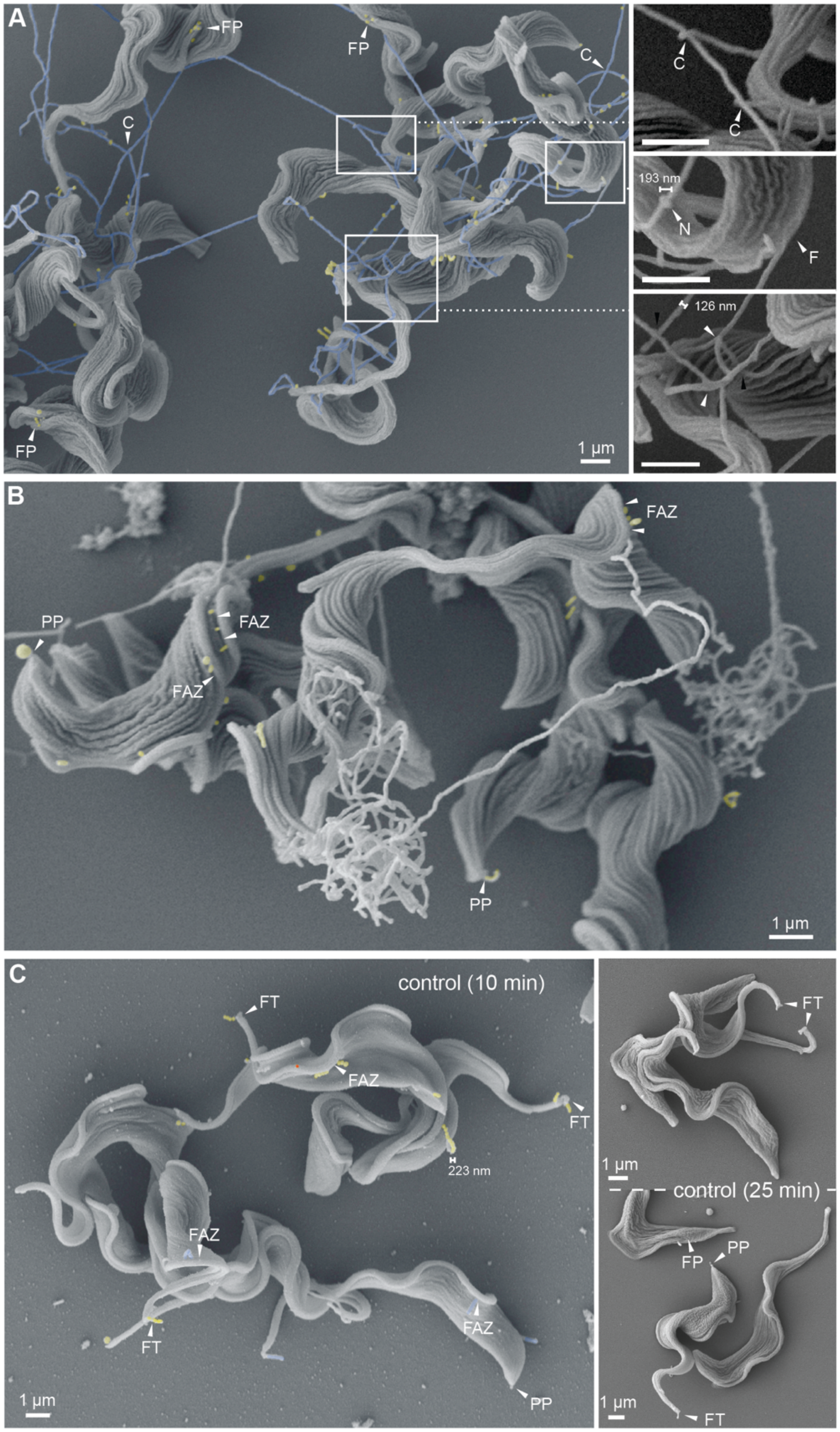

**Figure S10. Characteristics of Nanotube Intersections and Presence of Nanovesicles and Nanotubes in Untreated Cells, Related to Figure 6**

(A-C) SEM images of chemically fixed trypanosomes. FP, flagellar pocket; C, crossing; N, node; F, flagellum; FAZ, flagellar attachment zone; PP, posterior pole. (A-B) NB11$_{VSG2}$ treated VSG2 expressing cells. At 20 min post addition of Nb (A) the extended nanotubes (blue) reveal crossings, nodes and some fusion events. (B) Though long nanotubes are the obvious feature from 15 min post addition of Nb, even after 25 minutes nanovesicles (yellow) can still be seen originating from the FAZ and PP. (C) SEM images of control cells prepared in the same way as cells in (A) and (B), but without Nb treatment. Times in brackets indicate incubation time prior to fixing the cells. Though less abundant than in Nb treated cells, nanovesicles (yellow) and nanotubes (blue) of similar geometry, and originating from the FT, PP and FAZ, are also present in untreated cells.

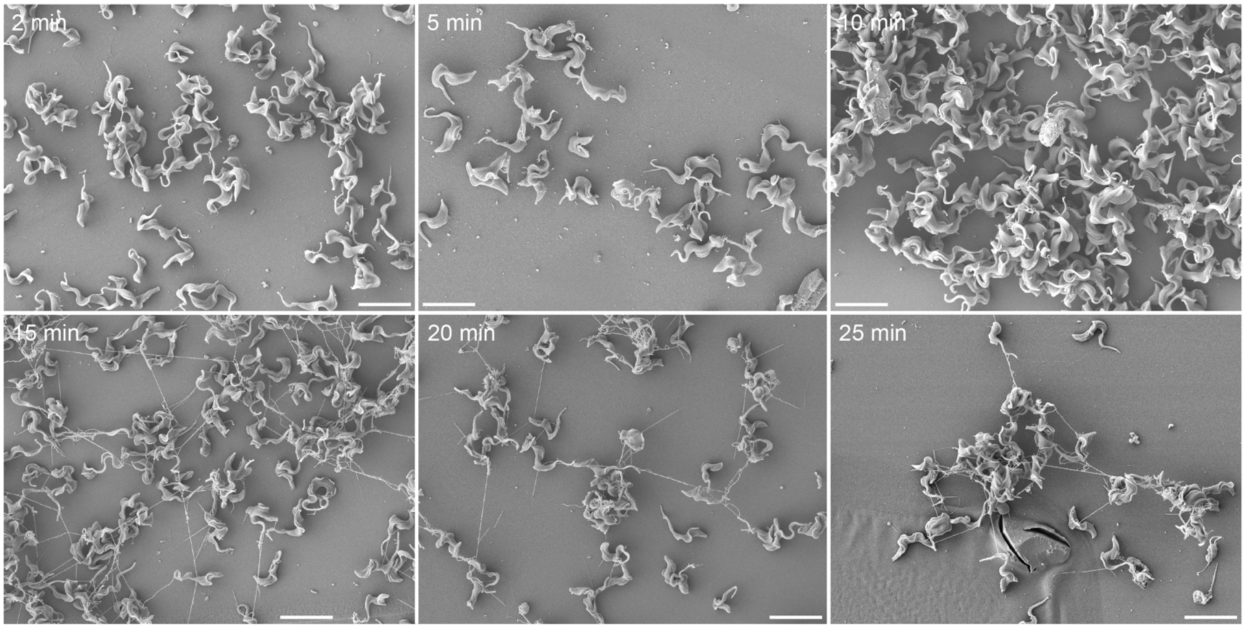

**Figure S11. Overview of SEM Images at Different Timepoints Following $NB11_{VSG2}$ of VSG2 Expressing Cells, Related to Figure 6**

Overview showing SEM images at 2, 5, 10, 15, 20 and 25 minutes post addition of $NB11_{VSG2}$ to VSG2 expressing cells. Nanovesicles and short nanotubes are visible at the earliest time points and become more elaborate over time. Scale bars, 10 μm.

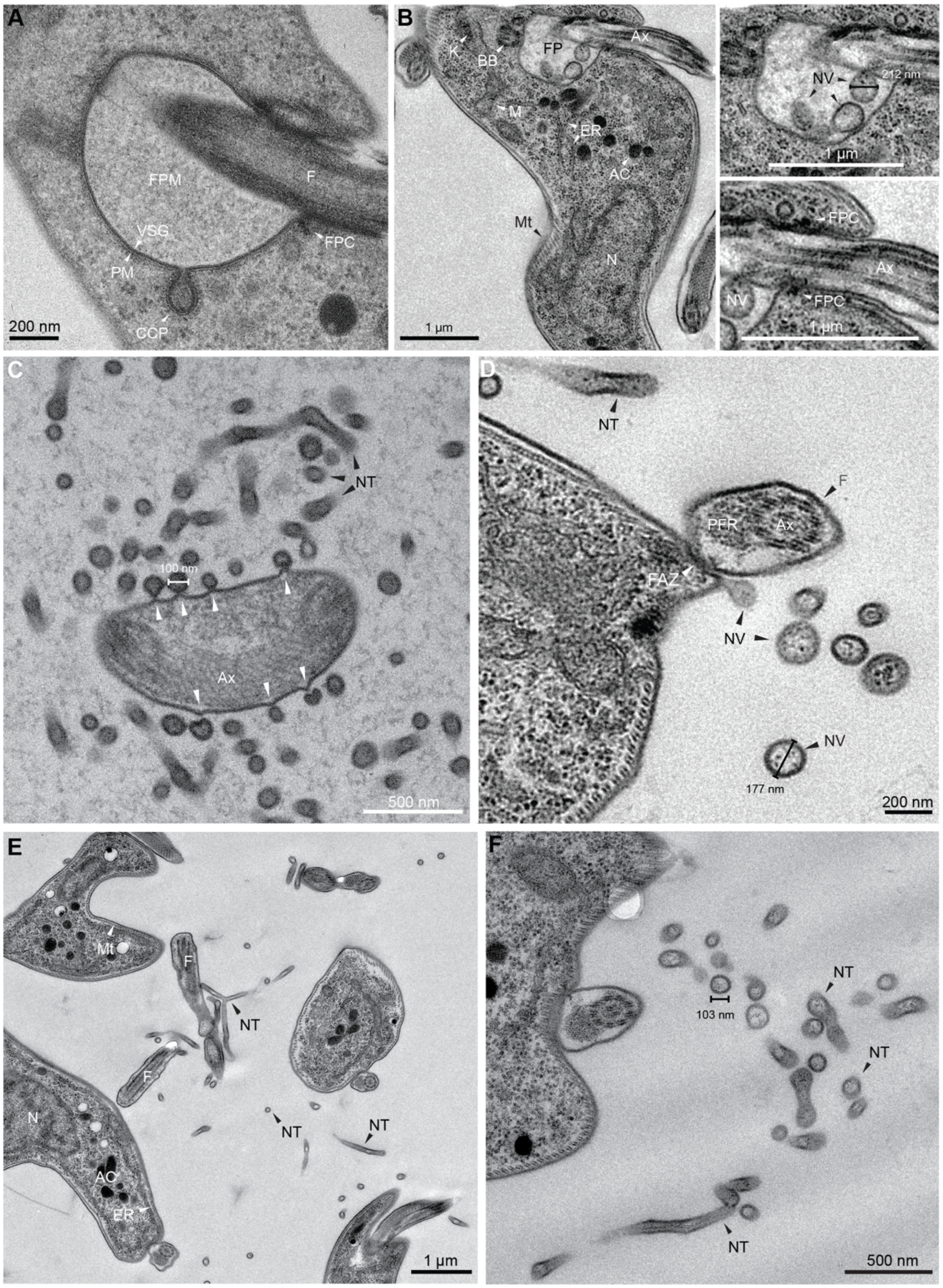

**Figure S12. TEM Images Reveal VSG coated Nanovesicle and Nanotubes in NB11$_{VSG2}$ Treated VSG2 Expressing Cells that Display an Unaltered Intracellular Architecture, Related to Figure 6**

(A-F) TEM images of NB11$_{VSG2}$ treated cells prepared by high pressure freezing (A) or chemical fixation (B-F). FPM, flagellar pocket matrix; F, flagellum; VSG, variant surface glycoprotein; PM, plasma membrane; CCP, clathrin coated pit; FPC, flagellar pocket collar; K, kinetoplast; BB, basal body; FP, flagellar pocket; Ax, Axoneme; M, mitochondrion; ER, endoplasmic reticulum; AC, acidocalcisome; N, nucleus; NV, nanovesicle; NT, nanotube. (A) Well preserved flagellar pocket showing the VSG coat on the pocket membrane, the flagellar pocket matrix and the flagellar pocket matrix. A clathrin coated pit, indicating an intact endocytic machinery is also captured. (B) Cross-section showing NVs in the flagellar pocket of an otherwise unremarkable cell displaying all the characteristic intracellular organelles. (C) Nanotubes lacking electro-dense structures appear to bud from the flagellum. Considerable deformations (white arrows) are visible on both the flagellar membrane and the nanotubes. (D) Nanotubes and nanovesicles containing electron-dense small structures, resembling ribosomes in the cytoplasm, most likely originate from the pellicular plasma membrane and not the flagellum. (E) Nanotubes were occasionally observed to have branchpoints. (F) Cross-sectioned nanotubes can be identified by their smaller diameter of around 100 nm compared to those of up to 200 nm in nanovesicles. Nanotubes can be curved and deformed.

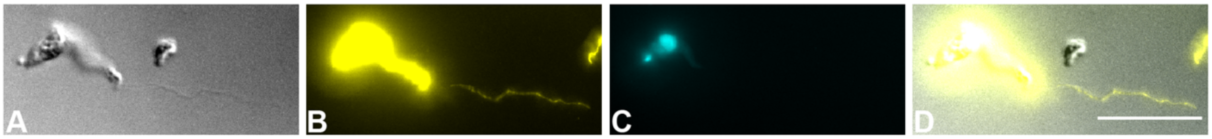

**Figure S13. Nanotubes are Covered with a VSG Coat, Related to Figure 6**

(A) DIC image of a trypanosome with a nanotube extending from the cell. (B) Immunofluorescence staining with anti-VSG2 antibody shows that, like the cell surface of trypanosomes, the nanotube is covered with VSG. (C) DAPI staining shows the single nucleus and kinetoplast. (D) Overlay of (A) and (B). Scale bar, 10 µm.

**Table S1: Crystallized Nb Sequences**

| Nanobody | Protein Sequence |
|---|---|
| $NB9_{VSG2}$ | QVQLQESGGGLVQAGGSLRLSCTTSGLTFSNYAFSWF<br>RQAPGEEREFVGAISWSGGRTDYADSVKGRFTISRDN<br>AKNTFYLQMNSLKTEDTAVYYCAADLLGEGSRRSEYE<br>YWGQGTQVTVSSAAAYPYDVPDYGS |
| $NB11_{VSG2}$ | QVQLQESGGGLVQAGGSLTLSCAVSGLTFSNYAMGWF<br>RQAPGKEREFVAAITWDGGNTYYTDSVKGRFTISRDN<br>AKNTVFLQMNSLKPEDTAVYYCAAKLLGSSRYELALA<br>GYDYWGQGTQVTVSSAAAYPYDVPDYGS |
| $NB14_{VSG2}$ | QVQLQESGGGLVQAGGSLRLSCEASGLTFSNYAMAWF<br>RQAPEKEREFVAGISWTGSRTYYADSVRGRFTTSRDG<br>HKNTVYLQMNDLKPEDTAVYLCAADLLGSGKDGTSVY<br>EYWGQGTQVTVSSAAAYPYDVPDYGS |
| $NB19_{VSG2}$ | QVQLQESGGGLVQAGGSLRLSCAASERTFSSLGM<br>GWFRQGPGKEREFAAAISWSGVSTYYADSVKGRF<br>TISRDNDKNTVYLQMNSLKPDDTAVYYCAATSSW<br>NDMALKSAGWYEYWGQGTQVTVSSAAAYPYDVPD<br>YGS |

### Table S2: Crystallographic Statistics

| | NB9$_{VSG2}$ | NB11$_{VSG2}$ | NB14$_{VSG2}$ | NB19$_{VSG2}$ |
|---|---|---|---|---|
| **Data Collection** | | | | |
| Beamline | SLS PXIII | Diamond I03 | Diamond I03 | Diamond I03 |
| Processing software | iMosflm | AutoProc | XIA2/DIALS | XIA2/DIALS |
| Wavelength (Å) | 1 | 0.9763 | 0.9763 | 0.9763 |
| Resolution (Å) | 47.782-1.499 (1.553-1.499) | 54.02-1.9 (1.986-1.9) | 74.24-1.3 (1.346-1.3) | 75.726-2.139 (2.216 -2.139) |
| Space group | P 21 21 21 | P 21 21 21 | P 21 21 21 | P 41 2 2 |
| Unit Cell *a*, b, *c* (Å) | 94.088 95.565 | 82.6438 97.7487 | 92.542 95.95 | 141.738 141.738 89.5825 |
| Unit Cell α, β, γ (°) | 123.721 90 90 90 | 104.506 90 90 90 | 124.337 90 90 90 | 90 90 90 |
| Total reflections | 11913750 (1027937) | 725832 (73662) | 3285113 (162714) | 1354843 (131277) |
| Unique reflections | 176929 (17100) | 67339 (6654) | 266024 (23312) | 50820 (4971) |
| Multiplicity | 67.3 (59.3) | 10.8 (11.1) | 12.3 (7.0) | 26.7 (26.4) |
| Completeness (%) | 99.18 (96.89) | 99.78 (99.61) | 98.27 (86.87) | 99.78 (98.82) |
| Mean *I/sigma(I)* | 51.47 (1.76) | 14.76 (1.43) | 17.72 (1.15) | 8.70 (1.46) |
| Wilson B-factor | 23.78 | 26.88 | 16.96 | 33.87 |
| *R-merge* | 0.6451 (3.386) | 0.2779 (2.104) | 0.05863 (1.281) | 0.3253 (2.75) |
| *R-meas* | 0.6501 (3.417) | 0.2926 (2.214) | 0.06107 (1.385) | 0.3316 (2.803) |
| *R-pim* | 0.07961 (0.4536) | 0.09019 (0.6773) | 0.01689 (0.5104) | 0.06378 (0.5407) |
| CC1/2 | 0.967 (0.594) | 0.948(0.187) | 1 (0.552) | 0.997 (0.591) |
| CC* | 0.992 (0.863) | 987 (0.562) | 1 (0.844) | 0.999 (0.862) |
| **Refinement** | | | | |
| Refinement reflections | 177002 (17100) | 67302 (6633) | 265900 (23312) | 50817 (4958) |
| R-free reflections | 8847 (853) | 3281 (336) | 13427 (1183) | 2448 (243) |
| R-work | 0.1641 (0.2489) | 0.2091 (0.3236) | 0.1512 (0.2880) | 0.2058 (0.2713) |
| R-free | 0.1916 (0.2856) | 0.2415 (0.3731) | 0.1723 (0.3091) | 0.2338 (0.3035) |
| CC (work) | 0.879 (0.615) | 0.929 (0.358) | 0.962 (0.811) | 0.956 (0.794) |
| CC (free) | 0.877 (0.636) | 0.941 (0.395) | 0.960 (0.789) | 0.932 (0.727) |
| No. of atoms | 7771 | 7798 | 8228 | 3872 |
| macromolecules | 6742 | 7122 | 7099 | 3638 |
| ligands | 104 | 78 | 154 | 39 |
| solvent | 925 | 598 | 975 | 195 |
| Protein residues | 939 | 964 | 932 | 484 |
| RMSD bonds (Å) | 0.005 | 0.007 | 0.01 | 0.008 |
| RMSD angles (degrees) | 0.78 | 0.90 | 1.18 | 0.96 |
| Ramachandran favored (%) | 99.04 | 96.63 | 96.93 | 97.07 |
| Ramachandran allowed (%) | 1.85 | 3.37 | 3.07 | 2.93 |
| Ramachandran outliers (%) | 0.11 | 0.00 | 0.00 | 0.00 |
| Rotamer outliers (%) | 0 | 0.98 | 0.14 | 0.27 |
| Clashscore | 0.91 | 2.40 | 0.91 | 1.93 |
| Average B-factor ($Å^2$) | 31.65 | 34.78 | 24.62 | 35.81 |
| macromolecules | 30.42 | 34.42 | 23.56 | 35.58 |
| ligands | 37.50 | 40.81 | 31.02 | 46.57 |
| solvent | 39.86 | 38.36 | 31.37 | 38.12 |
| TLS Groups | 0 | 0 | 0 | 0 |

Highest-resolution shell statistics are in parentheses. RMSD = root mean squared deviation.

**Table S3. *T. brucei* Survival is Dependent on the Concentration of $NB11_{VSG2}$**
Numerical values of the percentage of living cells during treatment with an approximate 0.85-, 1.7- and 3.4-fold molar excess of $NB11_{VSG2}$ to VSG monomer (3, 6 and 12 µg/mL $NB11_{VSG2}$ to 1.4 x $10^7$ cells/mL) over a time course of 8 h. All experiments together with an untreated control were performed in triplicate. Values represent the mean and standard deviation of all three experiments. Graphs for one replicate each shown in Figure 5C and Figure S8A.

| | living cells in % | | | |
|---|---|---|---|---|
| **time** | **0.85x** | **1.7x** | **3.4x** | **control** |
| **0 h** | 98.50±0.12 | 98.03±0.12 | 98.53±0.16 | 98.46±0.08 |
| **1 h** | 98.46±0.14 | 80.35±2.58 | 78.50±6.75 | 98.53±0.18 |
| **2 h** | 98.43±0.20 | 67.90±0.53 | 61.09±3.22 | 98.51±0.21 |
| **3 h** | 98.44±0.13 | 60.00±1.59 | 54.65±1.86 | 98.40±0.31 |
| **4 h** | 98.15±0.13 | 50.48±4.16 | 47.58±1.64 | 98.20±0.52 |
| **5 h** | 97.98±0.24 | 40.42±4.81 | 40.93±5.31 | 98.12±0.23 |
| **6 h** | 97.85±0.26 | 31.39±5.31 | 34.75±6.51 | 97.96±0.26 |
| **7 h** | 97.75±0.11 | 26.40±3.21 | 29.45±6.46 | 97.98±0.28 |
| **8 h** | 97.47±0.13 | 25.73±5.31 | 25.76±4.16 | 97.84±0.34 |

### Methods

- **Key Resources Table**

| REAGENT or RESOURCE | SOURCE | IDENTIFIER |
|---|---|---|
| **Antibodies** | | |
| Anti-6x-His | Abcam | ab1206 |
| Anti-a-Tubulin | Abcam | ab0474 |
| Anti-Rabbit | Biorad | 170-6515 |
| Rabbit anti-VSG221 (polyclonal serum) | raised against native VSG221 purified from *Trypanosoma brucei brucei* (own purification), custom made by BJ-Diagnostik Bio Science | Rb anti-VSG221, 14013, 1st bleed, 05.03.2014 (lab internal) |
| Goat anti-Rabbit Alexa Fluor 488 | Invitrogen | A11008 |
| **Bacterial and Virus Strains** | | |
| WK6 | ATCC | 47078 |
| DH5a | ThermoFisher | 18265017 |
| **Biological Samples** | | |
| **Chemicals, Peptides, and Recombinant Proteins** | | |
| Tryptone | Sigma-Aldrich | 91079-40-2 |
| IPTG | Roth | 2316.4 |

| | | |
|---|---|---|
| Glucose | ACROS | 410950010 |
| Trisma base | SIGMA | 77-86-1 |
| EDTA | Thermo Fisher | 17892 |
| Ni-NTA | Qiagen | 166017069 |
| Sucrose | SIGMA | 57-50-1 |
| Imidazole | ROTH | X998.4 |
| HMI-9 | PAN Biotech | so-15701 |
| FBS | gibco | 2024-02 |
| L-Cysteine | SERVA | 190022 |
| β-Mercaptoethanol | SIGMA | 102039442 |
| Yeast extract | GERBU | 8013-01-2 |
| Ampicillin | Sigma-Aldrich | BCBZ9179 |
| Magnesium chloride | Merck | 7786-30-3 |
| HEPES | Roth | 0195.3 |
| Sodium chloride | Fisher Chemical | 7647-14-5 |
| Zinc chloride | Merck | 7646-85-7 |
| Q-Sepharose | GE Healthcare | 17-0510-01 |
| LysC Protease | NEB | 10052276 |
| Dipotassium phosphate trihydrate | SUPELCO | a1351199024 |

| | | |
|---|---|---|
| FreeStyle 293 Expression media | Gibco | 2085258 |
| Monopotassium phosphate | SIGMA | 7778-77-0 |
| Sortase A | Internal production | - |
| AAGG-5-FAM | Biomatic | - |
| SDS | Panreac Applichem | 7U014009 |
| TBS | Merck | MFCD00132476 |
| Tween-20 | BIO RAD | 1610781 |
| Skimmed Milk Powder | SERVA | 68514-61-4 |
| Methylcellulose | Sigma-Aldrich | MO512 |
| Propidium iodide | Sigma-Aldrich | P4170 |
| Glutaraldehyde | Merck | 1.04239.0250 |
| Disodium hydrogen phosphate | Applichem | A4732.1000 |
| Acetone | Applichem | 131007.1611 |
| Gold-palladium | Baltic Praeparation | BP 2229 |
| Cacodylate buffer | Roth | 5169.2 |
| Osmium tetroxide | Electron Microscopy Sciences | 19130 |
| Uranyl acetate | Merck | 8473 |
| Ethanol | Th. Geyer | 2246.2500 |
| Propylene oxide | Sigma-Aldrich | 110205-2.5l |

| | | |
|---|---|---|
| **Epon 812** | **Serva** | **21045.02** |
| **Epon accelerator** | **Serva** | **36975.01** |
| **DDSA** | **Serva** | **20755.02** |
| **MNA** | **Serva** | **29452.03** |
| **Tannic Acid** | **Applichem** | **A3619.0250** |
| **Pioloform** | **Plano** | **R1275** |
| **Copper grids** | **Plano** | **G2500C** |
| **Lead (II) nitrate** | **Merck** | **1.07398.0100** |
| **tri-Sodium citrate dihydrate** | **Applichem** | **A2403.0500** |
| **Potassium chloride** | **Applichem** | **A3582.1000** |
| **BSA** | **Applichem** | **A1391.0100** |
| **Vectashield** | **Biozol** | **VEC-H-1000** |
| **DAPI** | **Applichem** | **A1001,0010** |
| **Magnesium sulfate** | **Serva** | **28311** |
| **Sodium dihydrogen phosphate** | **Applichem** | **A1373.1000** |
| **ATTO 488 NHS-ester** | **ATTO-TEC GmbH** | **AD488 31** |
| **Type A gelatin** | **Sigma-Aldrich** | **G18900-500G** |
| **Hellmanex** | **Hellma Analytics** | **9-307-011-4-507** |
| **PEG 1500** | **Sigma-Aldrich** | **225322-68-3** |

| | | |
|---|---|---|
| **ADA** | **Sigma-Aldrich** | **10202164** |
| **Citrate** | **Sigma Aldrich** | **102206486** |
| **Ammonium sulfate** | **Sigma Aldrich** | **BCBX0350** |
| **Magnesium sulfate** | **Merck** | **7487-88-9** |
| **PEG 400** | **Merck** | **25322-66-3** |
| **LB Broth** | **SIGMA** | **102123542** |
| **Critical Commercial Assays** | | |
| **FITC Conjugation Kit - Lightning** | **Abcam** | **ab102884** |
| **Pierce ECL** | **Thermo Fisher** | **32209** |
| **NEB HiFi DNA Assembly kit** | **New England Biolab** | **E5520S** |
| **Q5 High-Fidelity DNA Polymerase kit** | **New England Biolab** | **E0555S** |
| **BCA protein concentration kit** | **Pierce** | **23227** |
| **Deposited Data** | | |
| **VSG2(NB9VSG2)** | **This Paper** | **7AQX** |
| **VSG2(NB11VSG2)** | **This Paper** | **7AQY** |
| **VSG2(NB14VSG2)** | **This Paper** | **7AQZ** |
| **VSG2(NB19VSG2)** | **This Paper** | **7AR0** |
| **Experimental Models: Cell Lines** | | |
| **Experimental Models: Organisms/Strains** | | |

| | | |
|---|---|---|
| *Trypanosoma brucei brucei* 2T1 | **David Horn, University of Dundee** | **(Alsford et al., 2005)** |
| *Trypanosoma brucei brucei* Lister 427, clone 221 | **George Cross, Rockefeller University** | **http://tryps.rockefeller.edu/DocumentsGlobal/lineage_Lister427.pdf** |
| **Oligonucleotides** | | |
| **NB19_Y115A_rev** | **5'TGGCCCCATGCCTCATACCACCCTGCACTTTTAAGG-3'** | |
| **NB19_Y115A_fwd** | **5'TAT GAGGCATGGGGCCAGGGGACCCAGG-3'** | |
| **NB19_E114A_Y115A_fwd** | **5'GTGGTATGCGGCCTGGGGCCAGG-3'** | |
| **NB19_E114A_Y115A_rev** | **5'CCTGCACTTTTAAGGGCCATGTCG-3'** | |
| **NB9/11/14_universal_fwd** | **5'-GATGTGCAGCTGCAGGAGTCTGGRGGAGG 3'** | |
| **NB9/11/14_universal_rev** | **5'-CTAGTGCGGCCGCTGAGGAGACGGTGACCTGGG T-3'** | |
| **NB19_fwd** | **5'TCCCTGGACCCTGGCGGAACCAGCC-3'** | |
| **NB19_rev** | **5'GGCTGGTTCCGCCAGGGTCCAGGGA-3'** | |
| **cHC-Cam-IgG2_fwd** | **5'GGTCACCGTCTCCTCAGAGCCAAAGATTCCGCAG-3'** | |
| **cHC-Cam-IgG2_rev** | **5'-CCTGCAGCTGCACCTGACACTGAACCCCCTTCA-3'** | |
| **pHen6C-ER19_fwd** | **5'-GGAAAACCCTGGCGTTACCCAA-3'** | |

| | |
|---|---|
| pHen6C-ER19_rev | 5’-CAATCCAGCGGCTGCCG TAG-3’ |
| NB19_Y114A_fwd | 5’-TGCATACTGGGGCCAGG GGACCCAGGTCA-3’ |
| NB19_Y114A_rev | 5’-CTGGCCCCAGTATGCATAC CACCCTGCACTTTTAAG-3’ |
| NB11_L100A_fwd | 5’-GTAACGACTACTACCTAA GGCCTTCGCCGCAC AGTAATAC-3’ |
| NB11_L100A_rev | 5’-GTATTACTGTGCGGCG AAGGCCTTAAGTAGTAGTCGT TAC-3’ |
| NB11_100/101/103A_fwd | 5’-CCAAAGCCAGTTCGTAACG ACTAGCACCTGCGGCCTTCGC CGCACAGTAATACACGG-3’ |
| NB11_100/101/103A_fwd | 5’-CCGTGTATTACTGTGCGGC GAAGGCCGCAGGTGCTAGTC GTTACGAACTGGCTTTGG-3’ |
| NB11_L103A_fwd | 5’-CCAGTTCGTAACGACT AGCACCTAAGAGCTTCGCCG-3’ |
| NB11_L103A_rev | 5’-CGGCGAAGCTCTTAGGTGC TAGTCGTTACGAACTGG-3’ |
| pHen6c_G5_His | 5’-AGGTCACCGTCTCCTCAG GGGGTGGAGGCGGCCTGCCG AGCACCGGTGGCCATCATCAC CATCACCACTAATAGAATTCA CTGGCCGTC-3’ |
| pHen6c_G5_Ct_fwd | 5’-TAGAATTCACTGGCCGT CGTTTTACAAC-3’ |
| pHen6c-G5-Ct-rev | 5’- TGAGGAGACGGTGACCT GGGT-3’ |
| Nb_pHen6C_Srt_fwd | 5’ CAGGTGCAGCTGCA GGAGTC-3’ |
| Nb_pHen6C_Srt_rev | 5’- TGAGGAGACGGTGACCT GGGTCC-3’ |
| pHen6C_Srt_F1 | 5’- CCCAGGTCACCGTCTCC TCA-3’ |

| | |
|---|---|
| pHen6C_Srt_R1 | 5-<br>GACTCCTGCAGCTGCACCTG-<br>3’ |
| **Recombinant DNA** | |
| pHen6C with NB9 insert | This Study |
| pHen6C with NB11 insert | This Study |
| Phen6c with NB14 insert | This Study |
| Phen6c with NB19 insert | This Study |
| Phen6c with sortaggable NB9 insert | This Study |
| Phen6c with sortaggable NB11 insert | This Study |
| pcDNA3.1 with sortaggable heavy chain NB11 | This Study |
| Phen6c with sortaggable NB14 insert | This Study |
| Phen6c with sortaggable NB19 insert | This Study |
| Phen6c with NB11 100A mutant | This Study |
| Phen6c with NB11 103A mutant | This Study |
| Phen6c with NB11 100/101/103A mutant | This Study |
| Phen6c with NB19 114A mutant | This Study |
| Phen6c with NB19 115A mutant | This Study |
| Phen6c with NB11 114/115A mutant | This Study |
| Phen6c with ‘irrelevant NB’ BabA | This Study |

| **Software and Algorithms** | | |
|---|---|---|
| **COOT** | **(Emsley et al., 2010)** | **https://www2.mrc-lmb.cam.ac.uk/personal/pemsley/coot/** |
| **Phenix Software Suit** | **(Adams et al., 2010)** | **https://www.phenix-online.org** |
| **Chimera** | **(Pettersen et al., 2004)** | **https://www.cgl.ucsf.edu/chimera/** |
| **CCP4 Software Suit** | **(Collaborative Computational Project, 1994)** | **http://legacy.ccp4.ac.uk** |
| **iMosfilm** | **(Battye et al., 2011)** | **https://www.mrc-lmb.cam.ac.uk/harry/imosflm** |
| **AutoProc** | **(Vonrhein et al., 2011)** | **http://www.globalphasing.com/autoproc/** |
| **XIA2/DIALS** | **(Winter et al., 2013)** | **https://dials.github.io/index.html** |
| **Imaris x64 Version 9.5.1** | **Oxford Instruments** | **https://imaris.oxinst.com/** |
| **Prism Version 8.4.3** | **GraphPad** | **https://www.graphpad.com** |
| **Fiji (ImageJ v2.0.0)** | **(Rueden et al., 2017)** | **https://fiji.sc** |
| **CellQuest Pro 6.0** | **Becton Dickinson Bioscience** | |
| **Live Acquisition Software** | **Thermo Fisher Scientific** | |
| **Offline Analysis** | **Thermo Fisher Scientific** | |
| **Other** | | |
| **HiLoad 16/600 Superdex** | **GE Healthcare** | |

| | | |
|---|---|---|
| **24 well plate** | **Sarstedt** | **83.3922.500** |
| **PD10 columns** | **Thermo Fisher** | **29922** |
| **Slide-A-Lyzer** | **Thermo Fisher** | **P00090074** |
| **Cell culture flasks** | **Merck** | **C7481-50EA** |
| **Vivaspin 500** | **Sartorius** | **VS0191** |
| **Amicon stirred cell** | **Merck** | **UFSCO5001** |
| **Amicon filtration membrane** | **Merck** | **PCGC04310** |
| **24 Well Hanging Drop Vapor Diffusion Plate** | **Jena Bioscience** | **CPL-132** |

### RESOURCE AVAILABILITY

#### Lead Contact

Further information and requests for resources and reagents should be directed to and will be fulfilled by the Lead Contact, Nicola Jones.

#### Materials Availability

Plasmids generated from this study can be obtained directly from the Lead Contact.

#### Data and Code Availability

The following structure files are available from the Protein Databank: 7AQX, 7AQY, 7AQZ and 7AR0.

### EXPERIMENTAL MODEL AND SUBJECT DETAILS

The following Eukaryotic cell lines were used: FreeStyle 293 F cells, *Trypanosoma brucei brucei* 2T1 and *Trypanosoma brucei brucei* Lister 427, clone 221. The Freestyle 293 F cells were obtained from Thermo Fisher (Resource table). The *Trypanosoma brucei brucei* 2T1 were originally created in the laboratory of David Horn, University of Dundee, and *Trypanosoma brucei brucei* Lister 427, clone 221 were obtained from George Cross, Rockefeller University.

*Trypanosoma brucei brucei* 2T1 and Lister 427, clone 221 were grown in a humidified 37 °C incubator with 5 % $CO_2$ using HMI-9 medium (PAN Biotech) supplemented with 10 % FBS (Gibco), 1.5 mM L-cysteine (SERVA) and 0.2 mM β-mercaptoethanol (SIGMA). Cells were maintained below 5 x $10^5$ cells/mL and only grown to a density of 4 x $10^6$ cells/mL prior to harvesting. Freestyle 293 F cells were maintained in FreeStyle 293 Expression Medium (Thermo Fisher, 12338018). Cell were maintained below 1 x $10^6$ cells and only allowed to grow to higher density after transfection.

### METHOD DETAILS

#### Production of Nanobodies

Nanobodies (NBs) were produced by the VIB Nanobody Core (Vrije Universiteit Brussel). A llama was subcutaneously injected on days 0, 7, 14, 21, 28 and 35, each time with 150 µg of the N-terminal domain of VSG2 from *Trypanosoma brucei brucei*. The adjuvant used was Gerbu adjuvant P. On day 40, about 100 mL of anticoagulated blood was collected from the llama for lymphocyte preparation. A VHH library was constructed from the llama lymphocytes to screen for the presence of antigen-specific nanobodies. To this end, total RNA from peripheral blood lymphocytes was used as template for first strand cDNA synthesis with an oligo(dT) primer. Using this cDNA, the VHH encoding sequences were amplified by PCR, digested with PstI and NotI, and cloned into the PstI and NotI sites of the phagemid vector pMECS. The library consists of about 7 x $10^8$ independent transformants, with 100% of transformants harboring the vector with the right insert size (size of the DNA sequences coding for VHHs). The library was panned for 3 rounds in solution on inactivated, UV-irradiated *Trypanosoma brucei brucei* cells expressing

VSG2. Phages from the library were incubated with the *Trypanosoma brucei brucei* cells after which the cells were washed and the binding phages were eluted. The enrichment for antigen-specific phages was assessed after each round of panning by comparing the number of phagemid particles eluted from *Trypanosoma brucei brucei* cells (output) with the number of phagemid particles used for panning (input). The phage outputs for the 1st, 2nd and 3rd rounds were about $5 \times 10^4$, $5 \times 10^4$ and $1.5 \times 10^7$, respectively. The phage output increased about 30-fold in the 3rd round, as compared to the outputs from the 1st and 2nd rounds. The input phage was consistently near $10^{11}$. In total, 190 colonies (95 from round 2 and 95 from round 3) were randomly selected for sequencing. After removal of redundant sequences, there were 89 unique VHH sequences belonging to 46 different CDR3 groups (B-cell lineages). The crude periplasmic extracts from representatives of all CDR3 groups were tested by flow cytometry for their specificity for the target antigen, using inactivated fixed *Trypanosoma brucei brucei* cells. An irrelevant nanobody (a nanobody specific for *Helicobacter pylori* BabA antigen) served as negative control. A mouse anti-VSG2 mAb was used as positive control. Based on flow cytometry data and B-cell lineage data, 69 different positive nanobodies were identified, belonging to 30 different CDR3 groups (B-cell lineages). The nanobodies were sub-cloned into pHEN6c vector and expressed in WK6 cells. The PCR reaction was performed using Q5 High-Fidelity DNA Polymerase PCR kit (10 µL Q5 Reaction Buffer 5x, 1 µL 10 mM dNTPs, 2.5 µL Primers, 10 ng template DNA, 0.5 µL Q5 polymerase, 10 µL 5X Q5 High GC Enhancer and filled up to 50 µL with $H_2O$). Amplification of the NB insert by PCR was using pMECS plasmids harboring the nanobody gene as template and the reaction contained 30 cycles of PCR, each cycle consisting of 30 seconds at 94 °C, 30 seconds at 55 °C and 45 seconds at 72 °C, followed by 10 minutes extension at 72 °C at the end of PCR. A fragment of about 400 bp was amplified and can be visualized by 1 % agarose gel. Amplified insert was introduced to a pHEN6c backbone using standard ligation techniques following the Roche dephosphorylation and ligation kit instructions.

Expression was induced by overnight growth in TB medium at 28 °C with 1 mM IPTG addition at an OD600 of 0.6-0.8 (final OD600 after overnight induction was typically between 25 and 30). Cells were lysed by osmotic shock in TES buffer (0.2 M Tris pH 8.0, 0.5 mM EDTA, 0.5 sucrose) and the clarified supernatant incubated with Ni-NTA slurry (Qiagen, Germany). After washing with PBS, the protein was eluted with 0.5 M imidazole and then dialyzed against PBS.

**Production of Nanobody Mutants**

Nanobody mutants were produced using the Q5 High-Fidelity DNA Polymerase PCR kit (10 µL 5x Q5 Reaction Buffer, 1 µL 10 mM dNTPs, 2.5 µL Primers, 10 ng template DNA, 0.5 µL Q5 Polymerase, 10 µL 5X Q5 High GC Enhancer filled up to 50 µL with $H_2O$) with primers designed to create the desired mutations. PCR settings were 98 °C for initial denaturation (30 sec) followed by 35 cycles of 98 °C (10 sec), 72 °C (30 sec) and 72 °C (2 min). Final extension was 72 °C for 2 min. The PCR product was gel purified and 50 ng of the product was used in the NEBuilder HiFi DNA Assembly kit. After incubation for 1 h at 60 °C 2.5 µL were used for a standard transformation protocol. All clones were verified by DNA sequencing. Expression and purification of positive clones was performed as described before for wildtype nanobodies. Complex formation was performed as described before for the wildtype, but with less protein (1 mg each corresponding to 1:4 molar ratio).

**Production of Nanobody Mutants**

Nanobody mutants (NB11 L101A and NB19 E114A, Y115A, and E114/Y115A) were produced using the Q5 High-Fidelity DNA Polymerase PCR kit (10 µL 5x Q5 Reaction Buffer, 1 μL 10 mM dNTPs, 2.5 μL Primers, 10 ng template DNA, 0.5 μL Q5 Polymerase, 10 μL 5X Q5 High GC Enhancer filled up to 50 μL with $H_2O$) with primers designed to create the desired mutations. PCR settings were 98 °C for initial denaturation (30 sec) followed by 35 cycles of 98 °C (10 sec), 72 °C (30 sec) and 72 °C (2 min). Final extension was 72 °C for 2 min. The PCR product was gel purified and 50 ng of the product was used in the NEBuilder HiFi DNA Assembly kit. After incubation for 1 h at 60 °C 2.5 μL were used in standard transformation experiments. Nanobody mutants for NB11 L100A, L103A, and L100/L101/S103A were produced using the instructions of the QuickChange Lightning kit (Agilent) at a PCR reaction temperature of 68 °C. Successful reactions were used for standard transformation experiments. All clones were verified by DNA sequencing. Expression and purification of positive clones was performed as described before for wildtype nanobodies. Complex formation was performed as described before for the wildtype, but with less protein (1 mg each corresponding to a 1:4 molar ratio).

**Production of Sortaggable Nanobody**

Sortaggable Nbs were created by cloning a DNA fragment encoding *Streptococcus pyogenes* Sortase A recognition peptide (LPSTGG) flanked by a flexible penta-glycine linker and hexa-histidine tag to the NB-CTD sequence. A pHen6c vector was modified to harbor a penta-glycine and hexa-histidine tag using Q5 High-Fidelity Polymerase (NEB) and HiFi Assembly Mix as described above. Next, Nb sequences were subcloned into this new vector, using the Q5 High-Fidelity Polymerase (NEB) and HiFi Assembly Mix (NEB) as described above. Successful reactions were sent for sequencing and positive plasmids used in standard transformation experiments.

**Synthesis of a His-tagged VSG2-irrelevant Nanobody (NB-ER19)**

A nanobody generated against *Helicobacter pylori* adhesin BabA has been described before (Moonens et al., 2016, ''NB-ER19'') and was used as a negative control in biopanning of the M13 phage display VHH library on UV-irradiated *Trypanosoma brucei* cells expressing VSG2 by VIB. A codon-optimized DNA encoding NB-ER19 was synthesized and cloned into pHen6C plasmid using Q5 High-Fidelity DNA polymerase (NEB) as previously described.

The pHEN6C-ER19 plasmid encoding ER19 nanobody with a C-terminal His-tag was transformed into *E. coli* WK6 competent cells and purified after IPTG induction as described before.

**Purification of VSG2**

*Trypanosoma brucei brucei* expressing VSG2 were cultivated *in vitro* in HMI-9 media (formulated as described by Hirumi and Hirumi, 1989) by PAN Biotech without FBS, L-cysteine, or β-mercaptoethanol), supplemented with 10 % fetal calf serum (Gibco), 1.5 mM L-cysteine, and 0.2 mM β-mercaptoethanol. Cells were cultured at 37 °C with 5 % $CO_2$. VSG2 was purified according to established protocols (adapted from Cross, 1984). Briefly, cells were pelleted (20 min, 3200 x*g*, 4 °C) and then lysed in 0.2 mM ice-cold $ZnCl_2$. The lysis mixture was centrifuged for 10 min at 10000 x*g* at 4 °C (SN0) and the pellet containing the membrane material was resuspended in pre-warmed (42 °C) 20 mM HEPES buffer, pH 8.0 with 150 mM NaCl. Following a second centrifugation, supernatant containing VSG protein (SN1 + SN2) was loaded onto an anion-exchange column (Q-Sepharose Fast-Flow, GE Healthcare), which had been equilibrated with 20 mM HEPES buffer, pH 8.0 with 150 mM NaCl. The flow-through containing highly pure VSG was concentrated in an Amicon Stirred Cell and optionally shock frozen in liquid nitrogen and stored at -80 °C. For crystallization experiments the C-terminus of VSG2 was cut using 1 μg Endoproteinase LysC per 1 mg of VSG2. Digest was performed for 90 min at room temperature. A final step of purification involved gel filtration on a HiLoad 16/600 Superdex 200pg column (GE Healthcare) in 20 mM HEPES pH 8.0, 150 mM NaCl.

**Purification and Crystallization of VSG2-Nanobody Complexes**

Purified NTD VSG2 (in 20 mM HEPES pH8.0, 150 mM NaCl) and purified nanobody (in PBS) were mixed in a ratio of 1:4, resulting in a final buffer of 5 mM HEPES pH 8.0, 139.5 mM NaCl, 2 mM KCl, 6 mM $Na_2HPO_4$, and 1.5 mM $KH_2PO_4$. The VSG2-nanobody complex was further purified by gel filtration on a HiLoad 16/600 Superdex 200pg column in 10 mM Tris pH 8.0. VSG2-nanobody complexes were concentrated to 5-8 mg/mL. Crystals were grown by vapor diffusion using hanging drops formed from mixing a 1:1 volume ratio of the protein with an equilibration buffer consisting of 20 % PEG 1500, 0.1 M citrate (VSG2-NB9), 0.1 M imidazole, 19 % PEG 1500 (VSG2-NB11), 0.1 M citrate, 23 % PEG 1500 (VSG2-NB14) and 0.1 M ADA pH 6.5, 2 M ammonium sulfate, and 0.1 M magnesium sulfate (VSG2-NB19). Samples were cryo-protected by transfer of the crystals into a buffer of the crystallization solution augmented with 10-20 % PEG 400.

**Structure Determination of VSG2-Nanobody Complexes**

VSG2-nanobody datasets were collected at the Paul Scherrer Institut Villingen, (SLS) and Diamond Light Source. All structures were solved by molecular replacement with the PHENIX software suite ((Adams et al., 2010) (specifically, PHASER (McCoy et al., 2007)) using the PDB entries 1VSG and ChainB of 5lhr as the search models. Refinement with PHENIX coupled to cycles of manual rebuilding produced the final models of the complexes (Supplementary Table 2).

**Expression, Purification and Sortagging of Camelid VHH-IgG2 Heavy-chain Antibodies**

A DNA fragment containing the hinge region from camel IgG2, CH2, CH3, *Streptococcus pyogenes* Sortase A acceptor sequence (LPSTGG) and a His-tag was codon-optimized for *Homo sapiens* codon usage and synthesized as a clone in pcDNA3.1 plasmid. Next, the VHH domains were PCR amplified from pHen6c plasmids using Q5 DNA polymerase (NEB) as previously described using the primer NB-Cam-IgG2-fwd and NB-Cam-IgG2-rev. pcDNA3.1 plasmids containing Camelid heavy chain IgG2 were linearized by PCR using the primer cHc-Cam-IgG2-fwd and cHc-Cam-IgG2-rev as described before. Finally, the pcDNA3.1-IgG2 vector and the VHH amplicons were assembled to create pcDNA3.1-VHH-IgG2 expression plasmids, using HiFi DNA Assembly Master Mix as described before.

The pcDNA3.1 plasmid containing VHH-IgG2 heavy-chain antibody was transiently transfected into suspension cultures of FreeStyle™ 293-F cells in FreeStyle™ 293 Expression Medium using 293fectin™ Transfection Reagent as recommended by the manufacturer (Life Technologies). Culture supernatants were collected and filter sterilized. Purification was done using Ni-NTA agarose resin (Qiagen). The equilibrated resin was incubated with the supernatant and left gently shaking for 1 h at 4 °C. Afterwards the column was washed with 5 column volumes of 20 mM Tris pH 8.0, 30 mM imidazole and 0.3 M NaCl. Finally, the antibodies were eluted with 250 mM imidazole, 20 mM Tris pH 8.0, 30 mM imidazole and 0.3 M NaCl, followed by removal of imidazole by Slide-A-Lyzer™ dialysis cassette (10 kDa MWCO, Life Technologies) and concentration by Amicon-15 ultrafiltration unit (10 kDa MWCO, Millipore). Sortagging reactions were done using 2 mg of antibodies, 100 μM Streptococcus pyogenes Sortase A and 500 μM AAGG-5-FAM peptide (Biomatik, USA) in PBS overnight at 4 °C. Excess AAGG-5-FAM FAM-peptide was removed by a gel filtration column (GE Healthcare) as recommended by the manufacturer. Finally, Sortagged heavy chain IgG2 antibodies were concentrated using an Amicon 15 ultrafiltration unit (3 kDa MWCO, Millipore) and the concentrations were measured using BCA assay kit (Life Technologies) using a BSA standard curve.

**Sortagging of VSG2 Nanobodies**

Sortaggable versions of VSG2 nanobodies, expressed by pHen6C-Srt plasmids, were purified from WK6 cultures after IPTG induction, using a Ni-NTA agarose resin (Qiagen, Germany), similar to non-sortagged nanobodies. A peptide (AAGG-5-FAM) carrying a C-terminal 5-FAM was synthesized and purified by HPLC (Biomatik, USA). His-tagged Sortase A derived from Streptococcus pyogenes was produced in BL21-DE3 using pSpSortA-pET28a plasmid and purified as previously described (Pinger et al., 2017). Sortagging reactions were performed using 1 mg of nanobody, 75 μM Sortase A and 350 μM FAM-labeled peptide in PBS overnight at 4 °C. Excess FAM-peptide was removed by a gel filtration column (GE Healthcare). Finally, Sortagged nanobodies were concentrated using an Amicon 15 ultrafiltration unit (3 kDa MWCO, Millipore) and the concentrations were measured using BCA assay kit (Life Technologies) using a BSA standard curve.

**Flow Cytometry Analysis of Heavy-chain IgG2 Binding to the Native VSG2 Coat on *T. brucei***

2 x $10^6$ 2T1 cells expressing VSG2 were centrifuged at 6000 x*g* for 10 min at 4 °C and resuspended in 200 μL cold HMI-9 media and incubated with a 4-fold molar excess FAM-

sortagged reconstructed Camelid IgG2(NB11) antibody (relative to VSG2) and left at 4 °C for 10 min with gentle shaking. The unbound antibodies were washed away with a cycle of adding the volume to 2 mL and afterwards centrifuging at 6000 x*g* for 10 min at 4 °C. The washed cell pellet was resuspended in 200 µL cold HMI-9 medium for flow cytometry analysis (FACSCalibur, BD). Non-stained 2T1 cells were used as a control.

#### Flow Cytometry Analysis of Nanobodies Binding to the Native VSG2 Coat on *T. brucei*

2 x $10^6$ 2T1 cells expressing VSG2 were centrifuged at 6000 x*g* for 10 min and resuspended in 200 µL cold HMI-9 medium. The resuspended pellet was stained with 4-fold molar excess of $NB9_{VSG2}$, $NB11_{VSG2}$, $NB14_{VSG2}$, and $NB19_{VSG2}$, relative to VSG2, and left at 4 °C for 10 min with gentle shaking. The excess unbound nanobodies were washed away with a cycle of resuspending in 2 mL cold HMI-9 medium followed by centrifuging at 6000 x*g* for 10 min at 4 °C. The washed pellet was resuspended in 200 µL HMI-9 medium and incubated with anti-His antibody (1:1000) in cold HMI-9 medium at 4 °C for 10 min with gentle shaking. Samples were resuspended in 2 mL cold HMI-9 medium as an additional wash step and centrifuged at 6000 x*g*. Samples were resuspended in 200 µL cold HMI-9 medium prior to flow cytometry analysis. Controls were established by using non-treated parasites (2T1) and parasites (2T1) treated with anti-His antibody.

#### Flow Cytometry Analysis of FAM-sortagged Nanobodies Binding to the Native VSG2 Coat on *T. brucei*

Sortaggable versions of VSG2 nanobodies were purified as mentioned above. 2 x $10^6$ 2T1 cells expressing VSG2 were centrifuged at 6000 x*g* for 10 min and resuspended in 200 µL cold HMI-9 medium. The resuspended pellet was stained with 4-fold molar excess FAM-sortagged nanobodies at 4 °C for 10 min with gentle shaking. The excess nanobodies were washed away with a cycle of resuspending in 2 mL cold HMI-9 medium followed by centrifuging for 10 min at 6000 x*g*. The washed pellet was resuspended in 200 µL cold HMI-9 medium for flow cytometry analysis (FACSCalibur, BD). Non-stained 2T1 cells were used as a control.

**Flow Cytometry Analysis of NB11$_{VSG2}$ Binding to Different Native VSG Coats on *T. brucei***

2 x $10^6$ 2T1 cells expressing either VSG531, VSG615 or VSG1954 were centrifuged at 6000 *g* for 10 min at 4 °C and resuspended in 200 µL cold HMI-9 medium. The resuspended pellets were stained with 4-fold molar excess of NB11 relatively to VSG2 at 4 °C for 10 min with gentle shaking. The excess nanobodies were washed away with a cycle of resuspending in 2 mL cold HMI-9 medium followed by centrifuging at 6000 *g* for 10 min. The washed pellets were resuspended in 200 µL cold HMI-9 medium and incubated with anti-His antibody (1:1000) in cold HMI-9 medium at 4 °C for 10 min with gentle shaking. Samples were washed by a cycle of resuspending in 2 mL cold HMI-9 medium and centrifuging at 6000 *g* for 10 min. Samples were resuspended in 200 µL HMI-9 medium for flow cytometry analysis (FACSCalibur, BD). Non-stained 2T1 cells and 2T1 cells only incubated with anti-His antibody were used as a control.

**Flow Cytometry Analysis of FITS-conjugated NBs Binding to the Native VSG2 Coat on *T. brucei***

Nbs were conjugated using the FITC Conjugation Kit – Lightning Link from abcam (ab102844). 20 µg Nbs were mixed with 2 µL of the modifier reagent and afterwards incubated with 10 µg of the fluorescent material. Samples were left at room temperature for 3 h and the reaction was stopped using 2 µL of the quencher reagent. $10^6$ 2T1 cells expressing VSG2 were centrifuged at

6000 *g* for 10 min at 4 °C and stained with 4-fold molar excess FITC-conjugates (relative to VSG2) in cold HMI-9 medium at 4 °C for 10 min with gentle shaking. The excess NBs were washed away by resuspending in 2 mL cold HMI-9 medium and centrifuging for 10 min at 6000 *g* at 4 °C afterwards. The cells were resuspended in 200 μL HMI-9 medium for flow cytometry analysis (FACSCalibur, BD). Non-stained 2T1 cells were used as a control.

**Western Blot Analysis**

2 x $10^6$ 2T1 cells expressing VSG2 were centrifuged at 6000 *g* for 10 min and resuspended in 200 μL cold HMI-9 medium. The resuspended pellet was stained with 4-fold molar excess of NB9, NB11, NB14, NB19 and NB-ER19 (BabA), respectively, and left at 4 °C for 10 min with gentle shaking. The excess nanobodies were washed away with three cycles of resuspending in 2 mL cold HMI-9 medium and centrifuging at 6000 *g* for 10 min at 4 °C afterwards. The washed pellet was resuspended in 200 μL cold HMI-9 medium and Laemmli-Buffer was added to a final concentration of 1x.

Samples were run on SDS-PAGE (4-15 %) at 120 V for 50 minutes. Proteins were transferred to a Nitrocellulose membrane by applying 100 V for 75 minutes in the cold using standard Western blot protocols. Transferred membrane was blocked by incubating in 10 mL 3 % milk in TBS-T at 4 °C for 1 h. After washing, the membrane was incubated with primary antibody (anti-His (rabbit) 1:1000 and anti-α-Tubulin 1:3000 (rabbit)) in blocking solution overnight at 4 °C. Excess primary antibody was washed away with three cycles of washing with TBS-T. Secondary antibody (rabbit, 1:2000 for anti-His antibody, 1:5000 for anti-α-Tubulin antibody) was applied by incubating in blocking solution for 1 h at 4 °C. The membrane was then washed 3 times with TBS-T. Membrane was treated with ECL substrate and bands were visualized using the GelDoc system.

#### *T. brucei* Culturing for Motility and Toxicity Assays

All motility and subsequent toxicity, electron microscopy and immunofluorescence assays were performed on VSG2 expressing wildtype cells (Lister 427, clone 221) as preliminary experiments (not shown) found these to be more robust swimmers compared to other VSG2 expressing cell lines such as 2T1 or 13.90 cells. Trypanosomes were cultivated at 37 °C and 5 % $CO_2$ in HMI-9 cell culture medium containing 10 % heat-inactivated fetal calf serum and generally maintained below a density of 5 x $10^5$ cells/mL.

#### Tracking of Individual Cells in a Population

For analysis of cell trajectories, 1.4 x $10^6$ cells were harvested by centrifugation at 1500 x*g* for 10 min at room temperature and resuspended in 50 µL HMI-9 medium. An equal volume of MC-HMI-9 medium (HMI-9 medium containing 1.1 % methylcellulose (MO512, Sigma Aldrich, St. Louis, USA)) was added to reach a viscosity of 25 mPas and a cell density of 1.4 x $10^7$ cells/mL. The respective nanobodies were added to a final concentration of 6 µg/mL, resulting in an approx. 1.7-fold molar excess of nanobody to VSG2 monomer present on the cells. For data acquisition 3 µL of the cell suspension was placed on a slide and covered with a coverslip. In a pilot experiment recording of 30 min duration starting 5 min post addition of the nanobody were acquired with a stereomicroscope (Leica MZ16FA, Leica Microsystems, Mannheim, Germany) equipped with a Plan Apo 5x LWD objective (Leica Camera AG, Wetzlar, Germany), a zoom factor of 35x, and a CCD pco.1600 camera (PCO, Kelheim, Germany) employing an exposure time of 250 ms. In subsequent experiments videos of 125 s were acquired at 5, 15, 25 and 35 min post addition of the respective nanobody and the field of view was changed in between recordings to ensure adequate cell numbers for analysis. Trajectories were visualized and analyzed with the Imaris x64 software (Version 9.5.1, Bitplane AG, Zurich, Switzerland).

**High-speed Microscopy of Single Cells**

High-speed microscopy was performed to analyze effects on motility on a single cell level. Cells were prepared as for ensemble measurements described above but using a final cell density of 7 x $10^6$ cells/mL. High-speed videos were captured with an inverted fully automated DMI 6000B wide-field microscope (Leica Microsystems, Mannheim, Germany) equipped with a Leica 63x glycerol immersion objective (NA 1.3) (Leica Microsystems, Mannheim, Germany) and an sCMOS camera (PCO, Kelheim, Germany) using a frame-rate of 250 fps.

**Cell Viability Assay via Flow Cytometry**

For each preparation of cells for flow cytometry analysis, 1.4 x $10^7$ cells were harvested by centrifugation at 1500 x*g* for 10 min at room temperature. The cell pellet was washed with 4 mL of pre-warmed and filtered (pore size 0.22 µm) HMI-9 and centrifuged at 1500 x*g* for 5 min at room temperature. All but 1 mL of the supernatant was discarded to achieve a cell density of 1.4 x $10^7$ cells/mL. The cell suspension was transferred to a 24-well plate for incubation at 37 °C and 5 % $CO_2$ for the duration of the time course. Alongside a no nanobody control the required amount of nanobody (1-12 µg/mL for a 0.3- to 3.4-fold molar excess of nanobody to VSG2 monomer present on the cells) was added per well and subsequent gentle agitation of the plate ensured mixing with the cell suspension. At various time points during incubation, 50 µL of the cell suspension were removed and diluted with 250 µL of filtered HMI-9. Cells were then incubated for 10 min with 1 µg of propidium iodide (PI) which serves as a dead cell fluorescence marker. The ratio of fluorescent, dead cells to non-fluorescent living cells was analyzed with a FACSCalibur Flow Cytometer (Becton Dickinson Bioscience, Franklin Lakes, USA).

**Sample Preparation for Scanning Electron Microscopy (SEM)**

To prepare samples for SEM, 1.4 x $10^7$ cells were harvested by centrifugation at 1500 x*g* for 10 min at room temperature and resuspended in 750 µL HMI-9 medium. Cells were then treated with a 1.7-fold molar excess of $NB11_{VSG2}$ (6 µg) for the desired durations of time. For negative controls, cells were prepared and incubated as above with the omission of Nb addition. After incubation, 250 µL of 25 % glutaraldehyde were added and cells were fixed for 1 h at room temperature. The fixative was removed by washing once with Sørensen buffer (18.2 % of 60 mM $KH_2PO_4$, 81.8 % of 60 mM $Na_2HPO_4$, pH 7.4) with centrifugation at 1500 x*g* for 5 min at 4 °C. Following resuspension of the cells in fresh buffer they were spun onto polylysine coated coverslips at 1500 x*g* and 4 °C for 10 min and washed twice with Sørensen buffer. Coverslips with attached trypanosomes were transferred into critical point drying vessels and dehydrated at 4 °C for 5 min each in a series of 30 %, 50 %, 70 %, 90 %, 100 % (5x) acetone. This was followed by an overnight incubation in 100 % acetone and subsequent critical point drying (CPD) according to the manufacturer's instructions (Mulisch and Welsch, 2015). Coverslips were then attached to specialized racks which were connected by a thin layer of conductive silver. Finally, samples were vapor plated with gold-palladium for 150 s. A JSM-7500F Scanning Electron Microscope (JEOL, Tokyo, Japan) was used for imaging.

#### Chemical Fixation and Subsequent Epon Embedding for TEM

During chemical fixation, 6 x $10^7$ cells were harvested by centrifugation at 1500 x*g* for 10 min at room temperature and all except 900 µL supernatant was removed. Nanobody was added to an approximate 1.7-fold molar excess compared to VSG2 monomer present (24 µg) and incubated for the desired time. Negative controls were prepared in the same way except for omission of Nb addition. Following incubation, cells were fixed by adding 100 µL of 25 % glutaraldehyde and incubating for 1 h at room temperature. The samples were then washed 5 times with 1 mL 0.05 M cacodylate-buffer (pH 7.2). For this, samples were centrifuged at 1500 x*g* for 5 min, the supernatant was removed, and fresh buffer added while taking care not to resuspend the cell pellet. For a second fixation and for contrasting, cell pellets were incubated with 100 µL 2 % $OsO_4$ in 0.05 M cacodylate-buffer for 1 h at 4 °C. Samples were washed 5 times with pure $H_2O$ for 3 min each (centrifugation was omitted unless the pellet became dislodged). Subsequently, 1 mL 0.5 % uranyl acetate was added before overnight incubation at 4 °C. Cells were again washed 5 times with pure $H_2O$ for 3 min each. To dehydrate the samples, the pellets were incubated sequentially for 30 min each at 4 °C with 50 %, 70 %, 90 %, and finally 100 % EtOH. This was followed by two 30 min incubations with 100 % EtOH, and three 30 min incubations with propylene oxide at room temperature. Then, 50 % Epon in propylene oxide was added and incubated overnight as the first embedding step, followed by two incubation steps with 100 % Epon for 2 h each. Finally, Epon was exchanged one last time and polymerized at 60 °C for at least 48 h.

**High-pressure Freezing and Epon Embedding for Transmission Electron Microscopy (TEM)**

Generally, 2.8 x $10^7$ cells were harvested by centrifugation at 750 x*g* for 3 min at room temperature. The supernatant was reduced to a residual amount of 1 mL medium and 1 mL heat-inactivated fetal calf serum was added as a cryoprotectant. Nanobody was added to a 1.7-fold excess compared to VSG2 monomer present (12 µg total) and incubated for various durations of time. As a negative control, cells were prepared and incubated in exactly the same way but without the addition of nanobody. Following incubation, samples were centrifuged at 750 x*g* for 3 min at room temperature. All but around 200 µL of supernatant was removed and the cell pellet was resuspended carefully. The cell suspension was transferred into a PCR tube and centrifuged for 10 sec at room temperature and 2000 x*g* with subsequent removal of most of the supernatant. Cells were then transferred to the freezing chamber (specimen carrier type A (100 µm) and specimen carrier type B (0 µm)). High-pressure freezing was performed with an EM HPM100 (Leica Microsystems, Mannheim, Germany) at a freezing speed of > 20000 $Ks^{-1}$ and a pressure of > 2100 bar. Samples were stored in liquid nitrogen until embedding.

For Epon embedding, the frozen samples were transferred into an EM AFS2 automated freeze substitution system (Leica Microsystems, Mannheim, Germany) and incubated in 0.5 % (v/v) glutaraldehyde and 0.1 % (w/v) tannic acid in anhydrous acetone for 96 h at -90 °C. The solution was changed once after 24 h. After incubation, the samples were washed four times with anhydrous acetone at -90 °C for 1 h each and finally contrasted in 2 % (w/v) $OsO_4$ in anhydrous acetone for 28 h at -90 °C. Following this, the temperature was raised to -20 °C within 14 h, left at -20 °C for 16 h, and increased to 4 °C within 4 h. Then the samples were washed four times with anhydrous acetone at 4 °C over a period of 2-3 h. Finally the temperature was adjusted to 20 °C over a time of 1 h. Cell pellets were removed from the carriers and transferred into 50 % Epon in acetone. After 5 h the Epon was exchanged with 90 % Epon in acetone, and left overnight at 4 °C. The next day, samples were incubated three times with 100 % Epon for 2 h each at room temperature. Polymerization occurred at 60 °C for a minimum of 24 h.

**Preparation of Stained and Contrasted Sections for TEM Analysis**

An Ultra Jumbo Diamond Knife (Diatome AG, Nidau, Switzerland) was used to cut 60 nm serial sections of Epon embedded samples. Sections were placed on pioloform coated slotted copper grids and incubated for 10 min with 2 % uranyl acetate in ultrapure $H_2O$, followed by incubation for 5 min with Reynolds lead citrate. Images were acquired with a 200kV JEM-2100 (JEOL, Tokyo, Japan) transmission electron microscope equipped with a TemCam F416 4k x 4k camera (Tietz Video and Imaging Processing Systems, Gauting, Germany).

**Immunofluorescence Assays**

For immunofluorescence imaging, 3 x $10^6$ cells were harvested by centrifugation at 750 x*g* for 5 min at room temperature. The cells were washed by resuspending in 5 mL of serum-free HMI-9 and again centrifuging at 750 x*g* for 5 min at room temperature. Then, the cell pellet was resuspended in 200 µL serum-free HMI-9 to achieve a cell density of 1.4 x $10^7$ cells/mL. Nanobody was added in a 1.7-fold molar excess compared to VSG2 present in the sample followed by incubation for 15 min at room temperature. After incubation, the samples were brought to a final volume of 1 mL with serum-free HMI-9 and fixed for 30 min at room temperature with a final concentration of 4 % formaldehyde. Fixed cells were washed with TDB and pelleted by centrifugation at 750 x*g* for 5 min following which most of the supernatant was removed. Cells were resuspended in the residual supernatant and applied to polylysine coated slides and left to settle for 1 h. Slides were washed with PBS, incubated for 10 min with PBS/100 mM Tris and again washed with PBS before blocking for 1 h with 1 % BSA in PBS. Cells were then incubated for 1 h with a 1:1000 dilution of the primary rabbit anti-VSG221 antibody in 1 % BSA in PBS. After washing three times in PBS for 5 min each, cells were incubated for 1 h in the dark with a 1:500 dilution of the secondary goat anti-rabbit-Alexa 488 antibody in 1 % BSA in PBS. Slides were again washed three times for 5 min in PBS and mounted using Vectashield with DAPI prior to imaging. Images were captured with an inverted fully automated DMI 6000B wide-field microscope (Leica Microsystems, Mannheim, Germany) equipped with a 63x glycerol objective (NA 1.3) (Leica Microsystems, Mannheim, Germany) and a CCD camera (PCO, Kelheim, Germany.

**Fluorescence Recovery after Photobleaching (FRAP) Experiments**

A total of 1 x $10^7$ cells were harvested and washed three times with ice-cold TDB (5 mM KCl, 80 mM NaCl, 1 mM $MgSO_4$, 20 mM glucose, 20 mM $Na_2HPO_4$, 2 mM $NaH_2PO_4$, pH 7.6) and centrifuged at 1500 x*g* and 4 °C for 10 min. Following this, TDB was added to achieve a final cell density of 1 x $10^8$ cells/mL and trypanosomes were incubated with a final concentration of 10 µM ATTO 488 NHS-ester for 15 min on ice in the dark to label the cell surface. Cells were washed three times with 1 mL ice-cold TDB to remove all unbound dye. Following the final wash the labeled cells were resuspended in 20 µL TDB and $NB11_{VSG2}$ was added to a 1.7-fold molar excess over VSG2 monomer present (4.3 µg). For immobilization 3 µL of cell suspension were mixed with 5 µL of 10 % (w/v) type A gelatin (from porcine skin; Sigma-Aldrich) in PBS, pH 7.8, warmed to 37 °C between two cleaned coverslip and mounted in a temperature controlled sample holder which was cooled to 21 °C for data acquisition. Coverslips were cleaned by two 10 min sonication steps (37 kHz, 320 W, sweep mode) in 2 % Hellmanex followed by at least one further sonication step in ultra pure water. Coverslips were rinsed thoroughly in ultrapure water following each sonication step.

To avoid analysis of dead cells, data acquisition was restricted to 60 minutes following nanobody addition and care was taken to analyze only those cells that showed residual beating of the free flagellar tip in the otherwise immobilized cells.

To determine the diffusion coefficient and mobile fraction of VSG on living trypanosomes, line fluorescence recovery after photobleaching (line-FRAP) experiments were performed. For this, an inverted, fully automated wide-field microscope (FEI Munich GmbH, Germany), equipped with a Polychrome V monochromator (FEI Munich GmbH, Germany), a Yanus digital scan head (FEI Munich GmbH, Germany) for the control of 473 nm and 561 nm lasers (Cobolt Inc, Solna, Sweden), appropriate excitation and emission filters, a CCD camera (Sensicam, 6.45 µm/px; PCO, Kelheim, Germany) and a 60x oil immersion objective (NA 1.45) (Nikon, Tokyo, Japan) were used. Live Acquisition (FEI Munich GmbH, Germany) software was used for data acquisition. Data was analyzed using the Offline Analysis software packages (FEI Munich GmbH, Germany).

**Quantification And Statistical Analysis**

For the cell viability assay, the ratio of fluorescent, dead cells to non-fluorescent living cells was analyzed with a FACSCalibur Flow Cytometer (Becton Dickinson Bioscience, Franklin Lakes, USA). All experiments were performed in triplicate. FACS data for anti-his, non-VSG2 coats, FITC labelled material and FAM-labeled material was analyzed in FlowJo (10.5.3) and statistical measurements were performed using Prism 8. Data was obtained using a FACSCalibur Flow Cytometer (Becton Dickinson).

For FRAP experiments data was acquired at a constant temperature of 21 °C. In every experiment, 10 pre-bleaching images ($I_{pre}$) were acquired with 100 ms exposure time in a cycle of 500 ms. A line-shaped area ($I_{FRAP}(t)$) was then irreversibly bleached with a 20 ms laser pulse at 25 % laser intensity. The width of the bleached area ($2X$) was determined by analyzing the average distance between areas that were unaffected by bleaching. To monitor the fluorescence recovery in ($I_{FRAP}(t)$), 150 images were acquired after bleaching with 100 ms exposure time at intervals of 500 ms. Data were analyzed according to Phair et al. and Hartel et al. (Phair et al., 2004, Hartel et al., 2015).

In brief, at each time point the average fluorescence intensity of the bleached region of interest (ROI), the whole cell ($I_{fg}$) and the background ($I_{bg}$) were measured. A routine within the Offline Analysis Software (Thermo Fisher Scientific) used these data to perform a double normalization according to Phair with an exponential function then fitted to the normalized FRAP data. From this, the half-recovery time of fluorescence ($\tau$) and the ratio of mobile to immobile proteins ($MF$) was determined. Once $\tau$ and $X$ were known, the lateral diffusion coefficient $D$ was calculated from $D = \frac{X^2}{4 * \tau}$.
